# PathCLAST: Pathway-Augmented Contrastive Learning with Attention for Spatial Transcriptomics

**DOI:** 10.1101/2025.06.30.662247

**Authors:** Minho Noh, Sungkyung Lee, Sunghyun Kim, Sangsoo Lim

## Abstract

**Motivation:** Spatial transcriptomics provides high-resolution insights into tissue architecture and disease progression. While recent computational methods have advanced spatial domain identification, many focus primarily on gene expression alone, which may limit biological interpretability and underexploit complementary data such as histological images and known gene-pathway associations.

**Results:** We present PathCLAST (Pathway-Augmented Contrastive Learning with Attention for Spatial Transcriptomics), a novel framework that integrates gene expression, histopathological image features, and curated pathway graphs through a contrastive learning strategy. By embedding gene expression within biologically grounded pathway-level graphs and aligning them with histological features, PathCLAST enhances spatial domain resolution and provides interpretable attention scores over functional pathways. Across three benchmark datasets, PathCLAST consistently outperforms existing methods in clustering accuracy–achieving a 37% improvement over STAGATE on the IDC dataset and uncovers domain-specific signaling programs and spatial heterogeneity. Additional analyses of spatial autocorrelation and inter-domain crosstalk demonstrate its potential to reveal localized biological processes and tumor microenvironmental dynamics. Our code is available at the following link: https://github.com/sslim-aidrug/PathCLAST.

## 1 Introduction

Spatial transcriptomics enables the simultaneous measurement of gene expression and spatial location, offering unprecedented opportunities to study cellular heterogeneity, tissue organization, and disease progression *in situ* [1, 2, 3]. Recent advances in high-resolution technologies, including sequencing- and FISH-based methods [4], have led to exponential growth in spatial omics data, necessitating the development of robust computational tools.

Several computational methods have been developed to identify spatial domains and cell states [5], yet spatial transcriptomics data present unique challenges. These include high dimensionality, sparsity, and the fact that each spatial spot often contains transcripts from multiple cell types [6, 7]. Furthermore, the lack of direct cell-type annotations and the difficulty of modeling spatial dependencies between spots limit the resolution of tissue structure [8]. While recent graph-based deep learning methods - such as STAGATE [9], ConGI [6], and Hist2ST [8] - have improved spatial domain identification, they are often sensitive to data noise and may have limited ability to integrate multi-modal information such as histopathological context[10, 11].

Early spatial clustering methods such as BayesSpace [7] and Giotto [12] incorporated spatial priors (e.g., via Bayesian or HMRF frameworks) to improve spot classification. However, these statistical models generally assume spatial smoothness and may not effectively capture complex gene interactions or cell-type transitions [6, 13]. Deep learningbased approaches such as SEDR [14], CCST [15], and DeepST [16] introduced latent gene embeddings and variational inference to improve spatial inference, yet they still rely on proximity-based assumptions that may not hold in heterogeneous tumor environments [6, 9].

To improve robustness, STAGATE [9] applied graph attention networks over spatial graphs, using only transcriptomic data. This makes it vulnerable to missing or noisy expression features. In contrast, histopathological images have been shown to provide complementary biological context and can predict gene expression with reasonable accuracy [17]. Large-scale histological datasets offer additional structural and phenotypic cues [18, 19], motivating their integration with gene expression profiles. This has led to the development of multimodal frameworks that seek to combine image and transcriptomic features for more comprehensive tissue characterization.

Several methods incorporate image features, including stLearn [20] and SpaGCN [21], which extract morphological features to refine spatial graphs. However, these methods use image-derived distances only during graph construction, not during model training, and may propagate spurious relationships introduced by noisy features [6, 9]. More recent contrastive learning methods, such as ConST [22] and ConGI [6], attempt to align gene expression and image modalities by pulling together similar representations.

PathCLAST integrates curated biological pathways by constructing pathway-level graphs that guide model attention, enabling both robust representation learning and improved interpretability. Rather than relying solely on abstract latent embeddings, PathCLAST assigns attention scores to biologically grounded functional modules, revealing mechanistic insights into spatial domain organization and tumor progression.

In this study, we present PathCLAST, a pathway-aware contrastive learning framework for spatial transcriptomics. By encoding gene expression through curated pathway graphs, PathCLAST leverages prior biological knowledge to impose structural constraints on learned representations. This design enhances both robustness and interpretability while improving the alignment between histological and transcriptomic modalities. Furthermore, we introduce pathway-level data augmentation to support contrastive learning and uncover domain-specific biological signatures. Together, these contributions address current limitations in spatial omics modeling by enabling more accurate, interpretable, and biologically grounded identification of spatial domains.

## 2 Methods

The proposed PathCLAST (Pathway-Augmented Contrastive Learning with Attention for Spatial Transcriptomics) was decomposed into four main parts (Figure 1).

**Figure 1.**
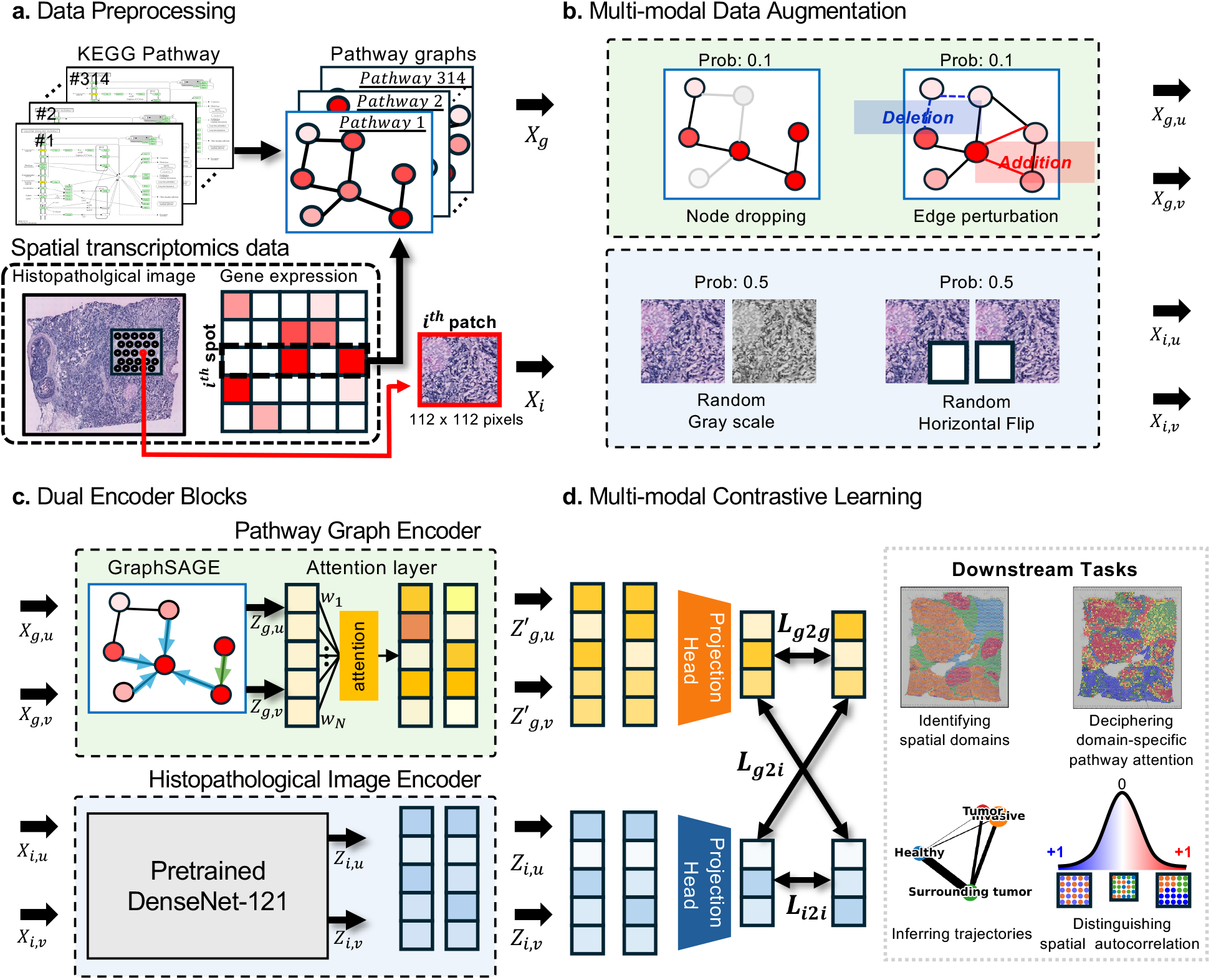
Architecture of the pathway-based spatial transcriptomics data augmentation contrastive learning model. PathCLAST jointly models spatial transcriptomics by integrating gene expression and histopathological image data through pathway-informed contrastive learning. (a) Gene expression profiles are transformed into biologically interpretable pathway graphs, while corresponding histological images are processed into spatially matched patches. (b) Both modalities undergo data-specific augmentationsgraph modifications for gene expression graphs and image transformations for image patchesto improve generalization. (c) These augmented data are encoded into low-dimensional embeddings via dedicated encoders. (d) Three contrastive objectives are used to align representations: within gene expression (*L*_*g*2*g*_), within histological images (*L*_*i*2*i*_), and across modalities (*L*_*g*2*i*_). After training, the model embeds raw data into a unified latent space, enabling downstream tasks such as domain-aware clustering, spatial visualization, and trajectory inference.

1. Data pre-processing and pathway-based graph construction for spatial transcriptomics data (Section: Data Preprocessing)
2. Data augmentation of both pathway graphs and image patches (Section: Multi-modal Data Augmentation)
3. Encoder blocks for pathway graphs and paired image patches (Section: Dual Encoder Blocks for Pathway Graph-Image Features)
4. Joint representation learning between modalities

(Section: Multi-modal Integration through Contrastive Learning)

### 2.1 Datasets

To benchmark our approach, we analyzed three publicly available spatial transcriptomics (ST) datasets:

#### HER2-Amplified Invasive Ductal Carcinoma (IDC)

We utilized the 10x Visium Human Breast Cancer dataset [23] (IDC), which consisted of 3,987 spots with a spot diameter of 55 *µm* and covering 36,601 genes. Based on histopathological annotations, spots were categorized into four primary morphotypes: ductal carcinoma *in situ* (DCIS), healthy tissue, invasive ductal carcinoma (IDC), and tumor edge regions with low malignant potential. Spot-level label information was provided in Supplementary Figure S2.

#### HER2-Positive Breast Tumor (Her2ST)

The HER2-positive breast tumor dataset [24] (Her2ST) was generated using spatial transcriptomics technology and included eight tissue slides with pathologist annotations. Each slide contained 300 − 700 spots with an individual diameter of 100 *µm*, covering over 15,000 genes. The dataset comprised three labels: normal tissue regions, ductal carcinoma *in situ* and invasive carcinoma. Slide C was excluded from this study due to its insufficient number of spots (fewer than 200).

#### Human Dorsolateral Prefrontal Cortex (DLPFC)

We also analyzed the 10x Visium Human DLPFC dataset [25] (DLPFC), which consisted of 12 tissue sections, each containing an average of 3,973 spots. This dataset included spatial gene expression data and histopathological images, enabling the study of spatial organization in the human prefrontal cortex.

### 2.2 Overview of PathCLAST

PathCLAST was a deep learning framework designed to identify spatial domains and decipher domain-specific pathway attention in spatial transcriptomics data by integrating pathway-based gene expression data with histopathological images (Figure 1). The framework consisted of the following components: (1) data preprocessing, (2) multi– modal data augmentation, (3) dual encoder blocks for pathway graph–image features, and (4) multi–modal integration through contrastive learning.

#### 2.2.1 Data Preprocessing

All three datasets consisted of paired gene expression profiles and histopathological images. Pathway graphs were first constructed using the KEGG pathway database [26]. Pathways lacking edges were removed, leaving 314 graphs covering 4,402 genes. Pathway graphs were generated with the KEGGgraph [27] library in R, and gene expression values were assigned as node features. For each spot, gene expression profiles were mapped to these pathway graphs, resulting in 314 pathway graphs collectively denoted as:

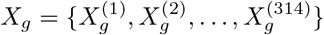

Histopathological images were extracted as 112 *×* 112 pixel patches from regions corresponding to each spot, denoted *X_i_* (Figure 1a).

#### 2.2.2 Multi-modal Data Augmentation

To improve model generalizability, both modalities underwent data augmentation. For each pathway graph in *X_g_*, we applied node dropping and edge perturbation, obtaining

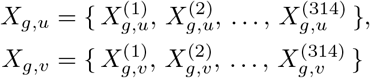

For image patches *X*_*i*_, we applied RandomGrayScale and RandomHorizontalFlip from the Torchvision package [28], and noise addition to obtain *X*_*i*,*u*_ and *X*_*i*,*v*_ (Figure 1b). Augmented data pairs were treated as positive samples for contrastive learning.

#### 2.2.3 Dual Encoder Blocks for Pathway Graph-Image Features

##### Pathway-Based Graph Encoder

Gene expression profiles were projected into low-dimensional embeddings using GraphSAGE [29] with an attention layer to learn pathway-level representations. The graph encoder, denoted as *f* ^graph^(·), processed each augmented pathway graph in *X*_*g*,*u*_ and *X*_*g*,*v*_, and produced latent features *Z*_*g*,*u*_ and *Z*_*g*,*v*_ for each pathway as follows:

##### Attention layer

The vanilla attention mechanism aggregated the node-level embeddings *Z*_*g*,*u*_ and *Z*_*g*,*v*_ into pathwaylevel representations 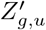 and 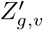. For each augmented view *m* ∈ *{u, v}* and node index *k* = 1, … , *n*, let 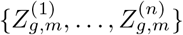 be the set of *n* node embeddings in the pathway graph. The attention score for node *k* was produced by the shared linear layer:

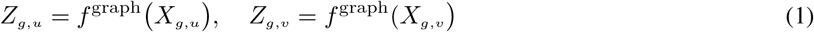

The attention weights were obtained by softmax normalization:

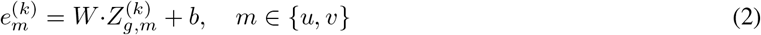

Finally, the pathway-level embeddings were then given by:

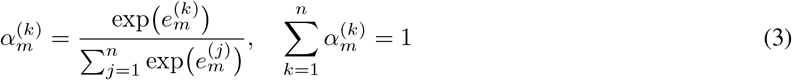

##### Image Encoder

For the image embeddings, DenseNet-121 [30] was adopted. Histopathological images were processed using DenseNet-121 convolutional neural network pre-trained on ImageNet [31]. The image encoder, denoted as *f* ^image^(·), processed the augmented image patches *X*_*i*,*u*_and *X*_*i*,*v*_as follows:

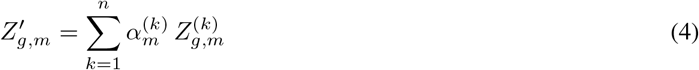

#### 2.2.4 Multi-modal Integration through Contrastive Learning

##### Projection Head

To align graph and image representations, projection head *f* ^pro^(·) projected embeddings into a joint latent space. The outputs were:

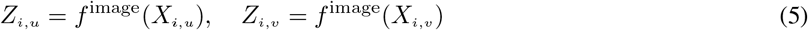

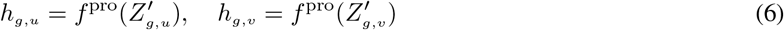

##### Multi-modal Contrastive Learning

We employed contrastive loss function based on the SimCLR framework [32]. To learn the joint representations between pathway graphs and images, we applied three contrastive loss termstwo within (graph-to-graph and image-to-image) and one between (graph-to-image). The contrastive learning between modalities aimed to pull paired low-dimensional representations *h*_*g*,*u*_and *h*_*i*,*u*_of pathway graphs and images together while contrasting those unmatching pairs apart. To better learn the characteristics of each modality, we also performed contrastive learning within each modality for pathway graphs and images, respectively.

A minibatch of *N* spots was randomly formed, and the contrastive prediction task was defined on paired spots derived from this minibatch, resulting in 2*N* paired spots. Negative spots were not explicitly constructed. Given a positive pair, the other 2(*N* − 1) spots in the minibatch were considered negative spots. Accordingly, the within- and between- contrastive loss functions for the positive pairs 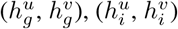 and 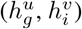 were defined as follows:

1. Graph-to-Graph Contrastive Loss (*L*_*g*2*g*_): Encourages alignment of augmented pathway graph embeddings.
2. Image-to-Image Contrastive Loss (*L*_*i*2*i*_): Aligns augmented image embeddings.
3. Graph-to-Image Contrastive Loss (*L*_*g*2*i*_): Aligns pathway graph embeddings with corresponding image embeddings.

The contrastive loss for a positive pair was defined as:

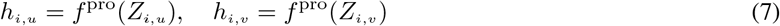

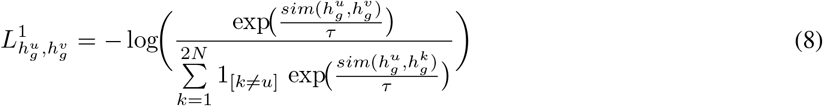

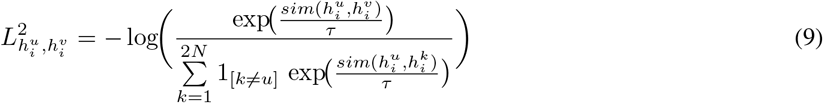

The indicator term **1**_[*k*≠*u*]_ was set to **1** when *k* differs from *u*, and *τ* was the temperature hyper-parameter. The similarity measure was defined as:

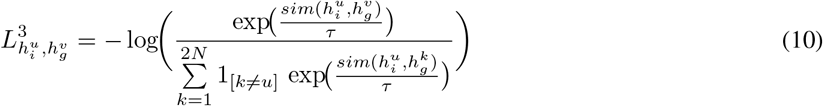

which corresponded to the cosine similarity between the two modality embeddings. The following equations calculated the average contrastive losses *L*_*g*2*g*_, *L*_*i*2*i*_, and *L*_*g*2*i*_ over all *N* pairs in the minibatch as follows:

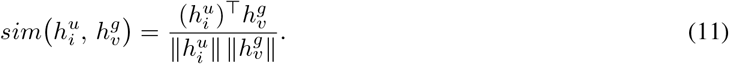

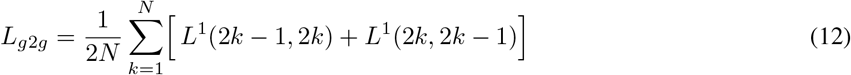

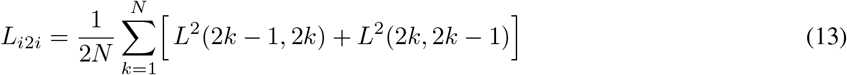

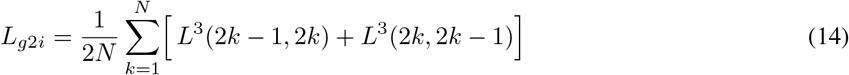

Finally, the total loss was computed as follows:

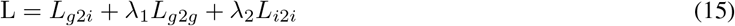

where *λ*_1_ and *λ*_2_ were hyper-parameters used to control the contributions of *L*_*g*2*g*_ and *L*_*i*2*i*_ to the final loss. *λ*_1_ = 0.3, *λ*_2_ = 0.2

### 2.3 Evaluation Criteria

The Adjusted Rand Index (ARI) was a measure of similarity between two clustering assignments, providing a robust evaluation of clustering performance [33]. ARI was insensitive to the number of clusters, allowing for a stable assessment of clustering quality. The ARI was defined as:

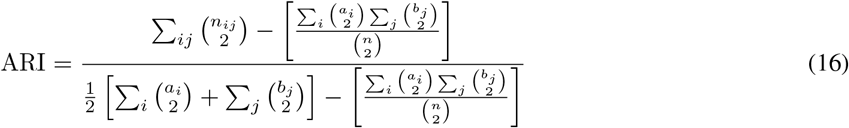

where:

- *n*: Total number of samples.
- *n_ij_*: Number of samples assigned to the *i*-th cluster in the predicted labels and the *j*-th cluster in the ground truth labels.
- *a_i_* = ∑*_j_n_ij_*: Total number of samples in the *i*-th predicted cluster.
- *b_j_* =∑*_i_n_ij_*: Total number of samples in the *j*-th ground truth cluster.

The ARI ranges from 0 to +1, where ARI = +1 indicates perfect agreement between the predicted and ground truth labels and ARI = 0 represents agreement equivalent to random chance.

Additionally, we employed Normalized Mutual Information (NMI), which measures the mutual dependence between predicted clusters and ground truth labels based on information theory [34]. The NMI was defined as:

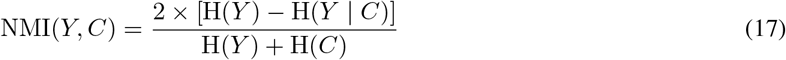

where:

- *Y* : Ground truth cluster
- *C*: Predicted cluster.
- *H*(*Y* ): Entropy of the ground truth cluster.
- *H*(*C*): Entropy of the predicted cluster.
- *H*(*Y* | *C*): Conditional entropy of the ground truth cluster given the predicted cluster.

Unlike ARI, NMI does not directly account for the number of clusters but measures the information shared between predicted clusters and ground truth clusters.

These indices ensure reliable comparison across datasets with varying cluster structures and sample sizes, making them ideal for evaluating spatial clustering performance in PathCLAST.

## 3 Results

### 3.1 PathCLAST Outperforms Previous Tools on Benchmark Datasets

PathCLAST was benchmarked on three publicly available spatial transcriptomics datasets: IDC, Her2ST, and DLPFC. Clustering performance was evaluated using ARI and NMI, measuring concordance with ground-truth spatial domains (Figure 2a). On the IDC dataset, PathCLAST achieved an ARI of 0.52, outperforming STAGATE (0.38) and ConGI (0.31) (Figure 2b). NMI scores reflected the ARI trend, reinforcing the robustness of PathCLASTs domain delineation. Performance gains were also observed on the Her2ST dataset, where PathCLAST attained a mean ARI of 0.46, substantially surpassing competing methods (Supplementary Figure S1).

**Figure 2.**
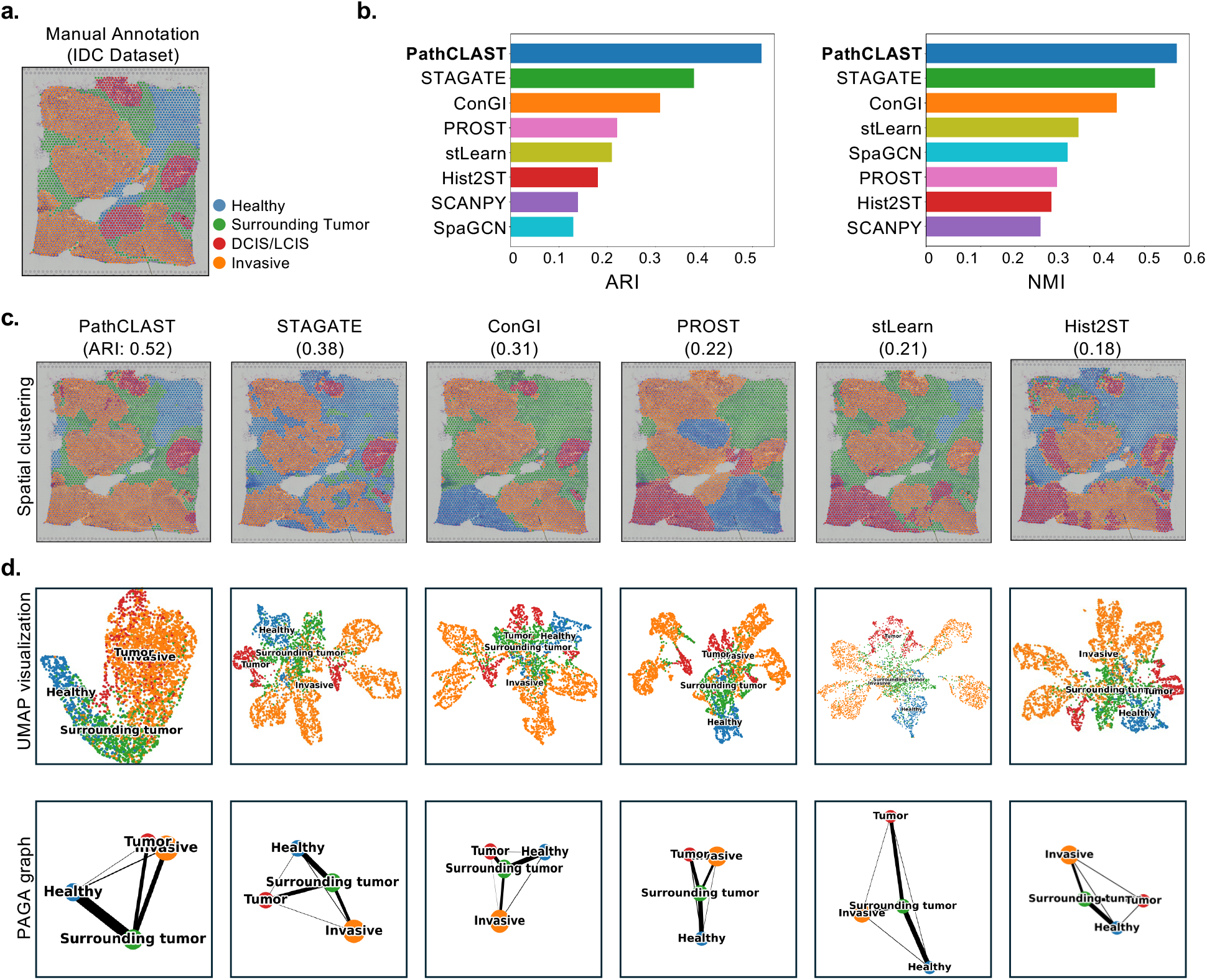
PathCLAST improves tumor identification in IDC tissue containing both invasive and DCIS/LCIS components. (a) Ground-truth segmentation of invasive tumor regions, DCIS/LCIS regions, and adjacent normal tissue in an IDC section. (b) Bar plots showing clustering accuracy (ARI and NMI) on the IDC dataset in terms of adjusted rand index (ARI) and normalized mutual information (NMI) scores for eight methods. (c) Predicted cluster assignments by PathCLAST, STAGATE, ConGI, PROST, stLearn, and Hist2ST in the IDC section. Other results are available in the Supplementary Figure S2. (d) UMAP visualizations and PAGA graphs generated by PathCLAST, STAGATE, ConGI, PROST, stLearn, and Hist2ST embeddings in the IDC section. For a complete comparison across all eight methods, see Supplementary Figure S3.

Spatial clustering maps (Figure 2c) revealed that most tools reliably identified healthy regions (blue) with strong consensus among methods, while notable discrepancies were observed in segmenting surrounding tumor regions (green). This was further quantified using contingency tables (Supplementary Figure S4). This suggests that PathCLAST effectively adapts to tissue heterogeneity, ensuring stable clustering across diverse spatial regions.

To further interrogate clustering quality, we visualized embeddings via UMAP and PAGA [35] (Figure 2d). PathCLAST demonstrated improved biological coherence and distinctly separates DCIS/LCIS, invasive, and healthy domains, outperforming STAGATE, ConGI, and other methods. These findings underscore the value of pathway-informed modeling in enhancing spatial domain resolution in transcriptomics data.

### 3.2 Pathway Attention Map Reveals Domain-specific Attention Profiles

To investigate the biological significance of the spatial domains identified by PathCLAST, we analyzed the pathway-level attention scores derived from our contrastive learning framework. Each spot-level gene expression was mapped to pathway-specific graphs, enabling PathCLAST to learn which pathways drive the clustering in different tissue regions.

A one-way ANOVA followed by Tukey’s HSD test (FDR *<* 0.05) was performed on spot-level spatial attention scores, which identified substantial differences in pathway attention among the four domains: invasive carcinoma, DCIS/L-CIS, surrounding tumor, and healthy tissue. Figure 3a presents Venn diagrams illustrating pathways significantly enriched in at least one domain. The full list of pathways, including statistical significance values and domain-specific classifications, is provided in Supplementary Table S3.

**Figure 3.**
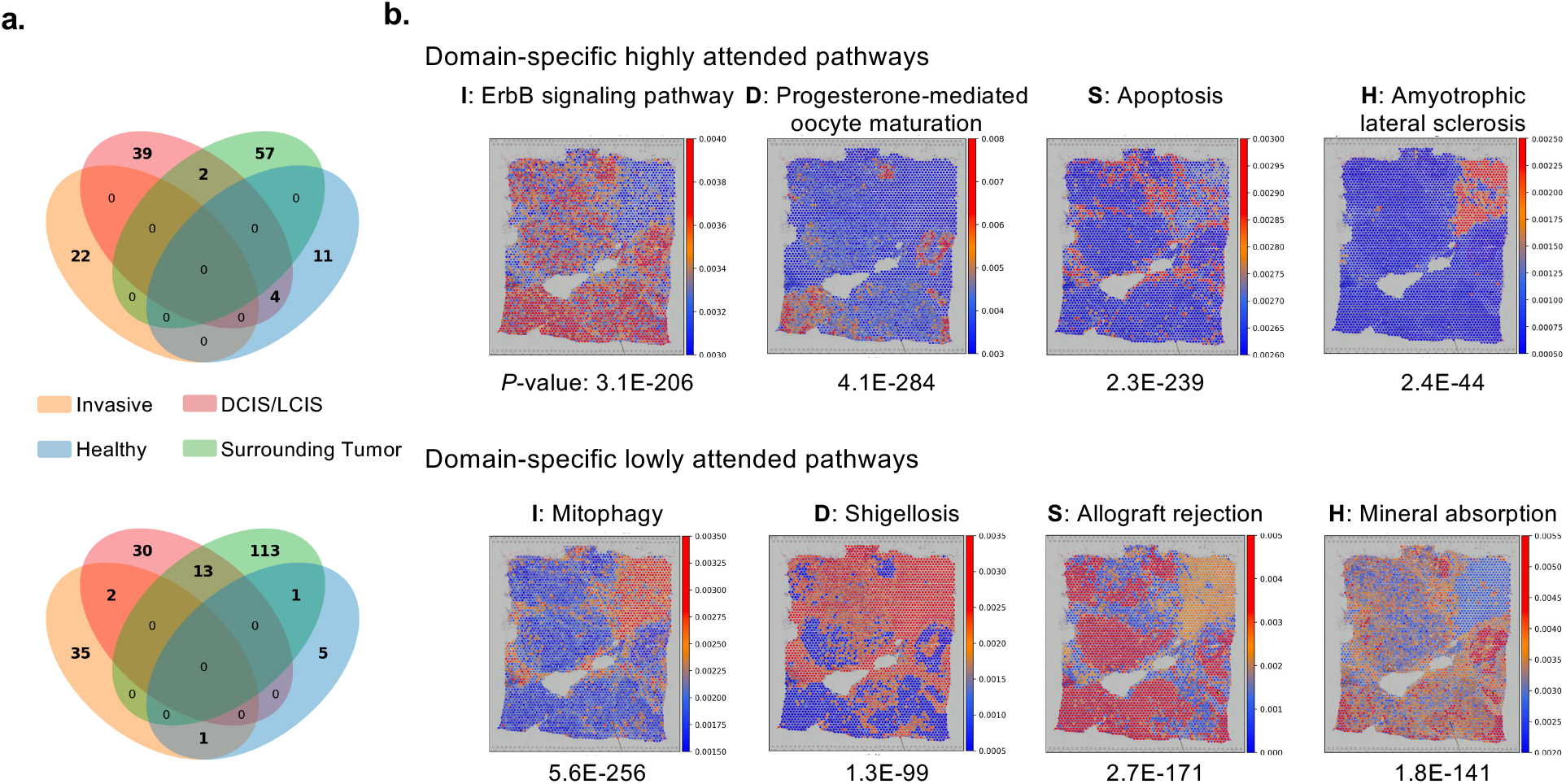
PathCLAST classifies domain-specific highly attended and lowly attended pathways using statistical analysis. (a) Pathway-level ANOVA across tissue labels (Invasive, DCIS/LCIS, Surrounding tumor, Healthy). Pathways that show significant differences then undergo post-hoc Tukey’s HSD analysis and are compared to the overall mean, allowing their classification as either domain-specific highly attended or lowly attended in each label. The results are visualized via two 4-way Venn diagramsone for domain-specific highly attended pathways (top) and one for domain-specific lowly attended pathways (bottom). (b) Pathways with the strongest domain-specific enrichment in ANOVA results (I: Invasive, D: DCIS/LCIS, S: Surrounding Tumor, H: Healthy). See Supplementary Table S3 for full results.

To interpret the spatial attention profiles learned by PathCLAST, we examined pathways assigned the highest and lowest attention weights across spatial domains (Figure 3b). High or low attention weights reflect the reliance of the model on specific pathways for spatial domain classification, not direct measurements of biological activity.

In invasive regions, PathCLAST prioritized the ErbB signaling pathway, consistent with its established role in driving epithelial malignancy [36, 37], while assigning low attention to mitophagy-related pathways, suggesting that mitochondrial quality control processes were less influential for classifying invasive tissues [38]. In DCIS/LCIS domains, high attention to the progesterone-mediated oocyte maturation pathway aligned with hormone-responsive proliferative features [39], whereas Shigellosis-related pathways were deprioritized, consistent with the limited relevance of pathogen response signatures in pre-invasive lesions [40]. Within the surrounding tumor microenvironment, apoptosis pathways received high attention, potentially reflecting tissue remodeling and immune-mediated clearance [41], while allograft rejection pathways were less informative [42]. In healthy tissue, PathCLAST assigned high attention to the Amyotrophic lateral sclerosis (ALS) pathway, potentially indicating the presence of homeostatic stress responses [43, 44], and low attention to mineral absorption pathways, suggesting nutrient transport features were not discriminative.

Collectively, 310 pathways were found to be significantly associated with domain classification (FDR *<* 0.05). Notably, Terpenoid backbone biosynthesis exhibited the strongest domain-specific enrichment (FDR-adjusted *p*-value = 1.4E − 303), highlighting the metabolic reprogramming characteristic of malignant tissues. The majority of highly attended pathways displayed domain-specific patterns, while lowly attended pathways helped suppress background biological signals, improving domain separability. For example, the high attention assigned to ErbB signaling across both tumor and surrounding regions hints at paracrine signaling interactions shaping the tumor microenvironment [45, 46, 47]. Similarly, domain-specific high attention to ALS pathways in healthy regions may reflect protective cellular mechanisms maintaining tissue integrity.

Overall, these domain-specific pathway signatures offer biological interpretability beyond purely data-driven clustering methods. Pathway-level attention scoring by PathCLAST enables the identification of candidate molecular processes underlying spatial organization and provides mechanistic insights critical for understanding disease progression.

### 3.3 Intra-Domain Heterogeneity of Pathway Profiles Along Tumor Progression

While domain-specific analysis reveals pathways distinguishing different spatial regions, tumors are intrinsically heterogeneous even within the same pathological domain. To capture this intra-domain diversity, Intra-domain heterogeneity was quantified by calculating the coefficient of variation (CV) of spot-level pathway attention scores within each labeled spatial domain (e.g., invasive carcinoma, ductal carcinoma *in situ*, healthy tissue).

Figure 4a categorizes intra-domain heterogeneity across six biological pathway groups. Among these, Metabolic pathways (MT) exhibited a distinctive pattern: invasive regions demonstrated uniformly high intra-domain heterogeneity, whereas DCIS/LCIS, surrounding tumor, and healthy regions displayed lower and more consistent heterogeneity across pathways within the same category. This consistent metabolic heterogeneity in invasive regions suggests pervasive spatial metabolic reprogramming, consistent with the known metabolic plasticity and adaptation observed in aggressive tumors [48, 49].

**Figure 4.**
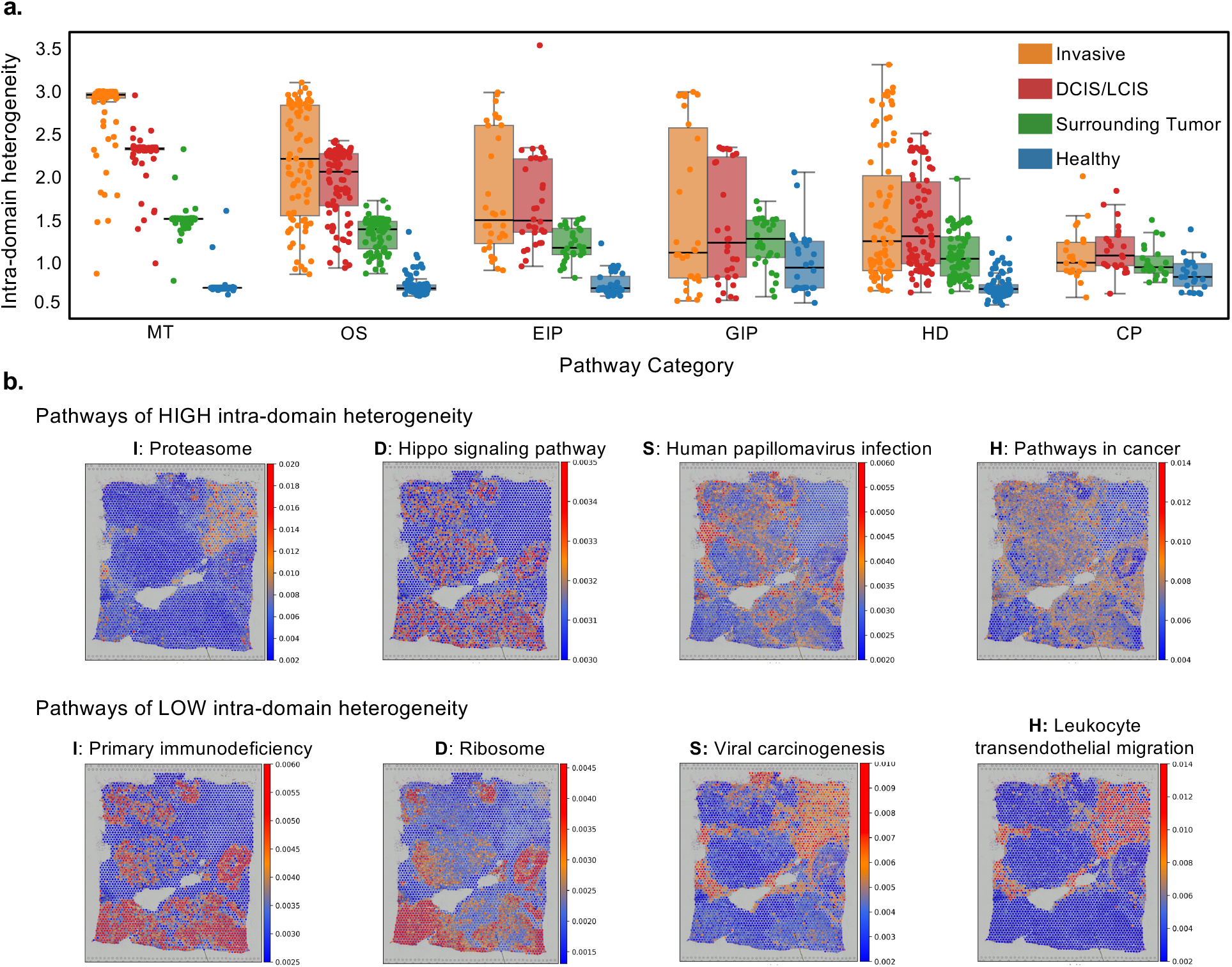
PathCLAST analyzes intra-domain heterogeneity of pathway categories across four tumor-related labels. (a) Boxplot of the computed intra-domain heterogeneity expressed as coefficient of variation for each pathway categorynamely, MT (Metabolism), OS (Organismal Systems), EIP (Environmental Information Processing), GIP (Genetic Information Processing), HD (Human Diseases), and CP (Cellular Processes). (b) Spatial visualization of pathways with high intra-domain heterogeneity (top) and pathways with low intra-domain heterogeneity (bottom).

In contrast, pathways related to Organismal Systems (OS), Environmental Information Processing (EIP), Genetic Information Processing (GIP), and Human Diseases (HD) exhibited a broader distribution of heterogeneity within invasive regions, indicating that proliferative, signaling, immune, and disease-related programs are spatially variable across tumor subregions. This variability was less pronounced in non-invasive compartments, where pathway attention scores were more tightly distributed [50, 51].

To further illustrate intra-domain heterogeneity patterns, Figure 4b presents spatial maps of selected pathways with either high or low intra-domain variability. Among pathways with high heterogeneity, Proteasome activity varied substantially within invasive regions, while the Hippo signaling pathway exhibited pronounced variability across DCIS/LCIS domains. Spatial heterogeneity of Human papillomavirus infection pathways in surrounding tumor regions suggests localized immune or viral mimicry dynamics, whereas Pathways in cancer showed unexpectedly high variability even within healthy tissue compartments. In contrast, pathways with low intra-domain heterogeneity included Primary immunodeficiency in invasive tumors, Ribosome pathways in DCIS/LCIS, Viral carcinogenesis in surrounding tumor regions, and Leukocyte transendothelial migration in healthy tissue. These findings indicate that while certain signaling and immune-modulatory pathways are spatially heterogeneous, fundamental immune functions and basic cellular processes remain relatively stable within given tissue domains.

In addition, high intra-domain heterogeneity was observed in the Thyroid cancer pathway (CV = 2.11) and Protein export pathway (CV = 2.03), particularly within invasive and surrounding tumor regions. Healthy tissue also exhibited localized variability, with pathways such as Amyotrophic lateral sclerosis (ALS) (CV = 1.56) showing elevated CV values, potentially reflecting baseline stress response heterogeneity even in normal tissues [43, 44]. The full pathway list of CV values is provided in Supplementary Table S4.

Spatial intra-domain heterogeneity of pathway attention scores, as captured by PathCLAST, suggests that histologically homogeneous tumor regions harbor molecularly distinct subpopulations with divergent proliferative, metabolic, and stress-response profiles. Such fine-grained diversity may underlie variable therapeutic responses, metastatic potential, and immune evasion within tumors [52, 53]. The ability to map these subdomain-specific pathway activations could facilitate more precise therapeutic targeting, especially in heterogeneous tumors where localized aggressive subclones may drive clinical progression. The ability of PathCLAST to spotlight these varying pathway activities at sub-domain resolution suggests its potential for informing personalized therapeutic strategiesfor example, by targeting subregions with elevated ErbB signaling pathway activity that may drive invasiveness or metastasis [36, 47].

### 3.4 Autocorrelation Analysis Reveals Spatial Dependence of Tumor Progression

We performed a spatial autocorrelation analysis to investigate domain-specific pathway activity across tumor, invasive, and healthy tissue regions identified by PathCLAST (Supplementary Figure S9). The analysis pipeline involved constructing domain-specific spatial networks, clustering pathways using the Markov Cluster Algorithm (MCL) [54], and quantifying the degree of spatial organization using Moran*^′^*s I statistic, where values range from -1 (indicating spatial dispersion) to +1 (indicating strong spatial clustering) [55].

First, MCL clustering revealed distinct intra-domain subdivisions across invasive, DCIS/LCIS, surrounding tumor, and healthy tissues (Figure 5b). Notably, invasive clusters (I1I4) generally showed higher Moran*^′^*s I values compared to healthy clusters (H1H3), indicating stronger spatial structuring of pathway activities within malignant compartments.

**Figure 5.**
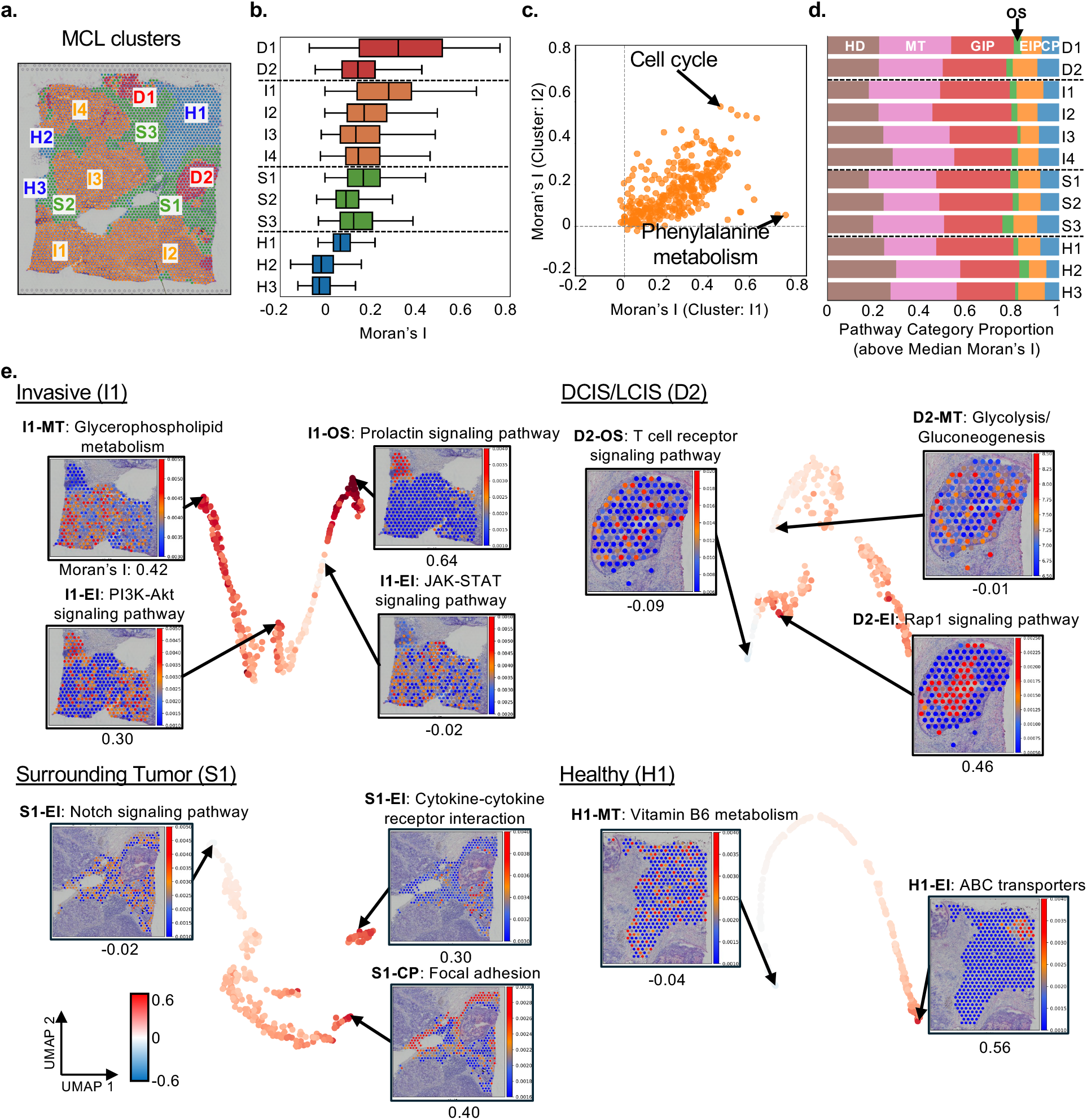
PathCLAST analyzes the spatial autocorrelation of pathway attention among sub-domains. (a) For each domain, MCL clustering is performed to further subdivide the domain. (b) After MCL clustering, pathway attention weights within each cluster are used to calculate Moran*^′^*s I, and the resulting distributions are visualized using boxplots. (c) The correlation of Moran*^′^*s I between two clusters (I1, I2) is presented as a scatterplot, with pathways corresponding to high data point values indicated. (d) A bar plot displays the proportion of pathway categories for pathways with Moran*^′^*s I values above the cluster median. (e) The computed Moran*^′^*s I values are visualized using UMAP, with each data point representing the Moran*^′^*s I value for a pathway. In addition, a spatial visualization of pathway attention weghts–filtered by cluster–is provided. A detailed schematic of the entire workflow is available in Supplementary Figure S9.

Correlation analysis of Moran*^′^*s I values between different clusters further demonstrated that spatial patterns of pathway activation were conserved across multiple tumor compartments (Figure 5c). For example, pathways related to cell cycle regulation and pyrimidine metabolism exhibited consistently high Moran*^′^*s I values across independent invasive clusters, suggesting conserved spatially restricted proliferation and metabolic activity during tumor progression.

At the functional category level, pathways assigned to Organismal Systems (OS), Environmental Information Processing (EIP), and Metabolism (MT) were predominant among pathways with Moran*^′^*s I values above the cluster median (Figure 5d). This indicates that spatial organization within tumors is particularly enriched for proliferative signaling, immune communication, and metabolic reprogramming pathways.

To illustrate these findings, we visualized the spatial distributions of selected high-and low-autocorrelation pathways (Figure 5e). Among pathways exhibiting strong clustering, the Prolactin signaling pathway (Moran*^′^*s I = 0.64) and the Rap1 signaling pathway (Moran*^′^*s I = 0.46) displayed regionally localized activation within invasive and DCIS/LCIS domains, respectively. These localized patterns suggest that spatially constrained activation of growth and adhesion pathways may underpin intratumoral niche formation and local progression. In contrast, pathways such as the T cell receptor signaling pathway (Moran*^′^*s I = -0.09) and Vitamin B6 metabolism (Moran*^′^*s I = -0.04) were more spatially dispersed, indicating broader tissue-level engagement of immune and metabolic programs.

Together, these analyses reveal that PathCLAST not only detects domain-specific pathway enrichments but also resolves the spatial autocorrelation of biological processes within tumors. The ability to map spatially organized versus spatially diffuse pathway activities offers new insights into how tumors compartmentalize key signaling, metabolic, and immune processes during progression and how microenvironmental heterogeneity may shape therapeutic responses.

The spatially localized activation of the Prolactin signaling pathway in invasive domains may reflect its known role in promoting tumor cell proliferation, angiogenesis, and survival in the breast tumor microenvironment [56, 57, 58]. Likewise, the regional clustering of Rap1 signaling pathway activity in DCIS/LCIS regions is consistent with its established function in regulating cell adhesion, polarity, and junctional integrity, processes that are critical during early tumor progression and local invasion [59, 60]. These spatially organized signaling programs highlight how tumors compartmentalize key molecular activities to support progression and adaptation to local microenvironmental cues.

DCIS/LCIS regions exhibited strong clustering, with the Rap1 signaling pathway (D2-EI) showing pronounced clustering, reflecting localized activity associated with DCIS/LCIS microenvironments. In invasive regions, the Prolactin signaling pathway (I1-OS) and Glycerophospholipid metabolism (I1-MT) demonstrated high Moran*^′^*s I values, indicative of spatially localized activation, whereas pathways such as the JAK-STAT signaling pathway (I1-EI) displayed broader distributions, suggesting spatial dispersion. In healthy regions, ABC transporters (H1-EI) exhibited strong clustering, while pathways like Vitamin B6 metabolism (H1-MT) showed a random spatial organization. These results underscore the ability of PathCLAST to uncover the spatial dynamics of pathways in tissue microenvironments, providing biologically meaningful insights into domain-specific activity and organization.

### 3.5 Domain-level Pathway Crosstalk Reveals Spatially Localized Characteristics of Cancer Progression

To investigate how spatially distinct regions coordinate biological processes during tumor progression, we performed a domain-level crosstalk analysis by computing Cross-Moran*^′^*s I between pathway attention scores across different tissue compartments (Figure 6a). Cross-Moran*^′^*s I quantifies the spatial similarity of pathway activities between two domains, revealing how molecular programs interact across tumor boundaries. The resulting network graph showed strong cross-domain interactions between invasive carcinoma, DCIS/LCIS, and surrounding tumor regions, with fewer high-strength connections involving healthy tissue. Particularly, invasive regions exhibited extensive crosstalk with both adjacent tumor edge (surrounding tumor) and DCIS/LCIS compartments, highlighting the interconnected nature of malignant progression.

**Figure 6.**
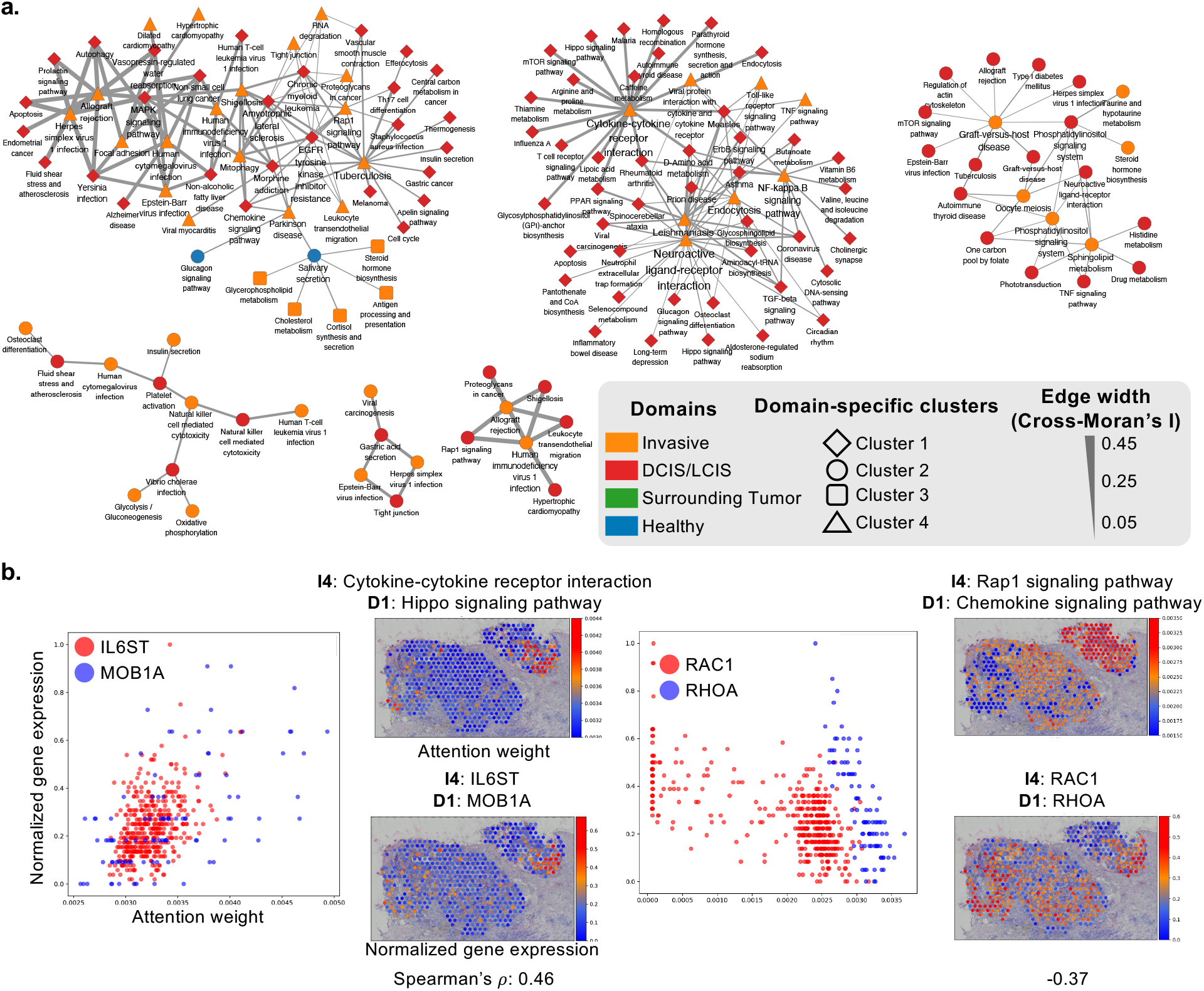
PathCLAST captures spatially resolved pathway crosstalk between domains. (a) A network of connected components is constructed by computing Cross-Moran*^′^*s I for pairs of domains, with edges representing the Cross-Moran*^′^*s I values. (b) Pathway attention weights for pathways exhibiting crosstalk and the corresponding normalized gene expression data for genes within each pathway are presented through spatial visualization. Additionally, scatter plots depict the correlation between pathway attention weights and normalized gene expression levels.

To validate the functional relevance of these spatial crosstalk patterns, we examined pathways with high Cross-Moran’s I scores (Figure 6b). Notably, the Hippo signaling pathway (highly active in DCIS/LCIS, D1) and the Cytokinecytokine receptor interaction pathway (invasive region, I4) exhibited strong crosstalk. This is biologically plausible given that the Hippo pathway integrates mechanical and inflammatory cues to regulate tumor growth and metastasis [61, 62], and Cytokine-cytokine receptor interaction pathway modulates Hippo signaling pathway activity through NF-*κ*B-mediated inflammatory responses [63]. Similarly, crosstalk between the Rap1 signaling pathway (I4) and the

Chemokine signaling pathway (D1) was observed, reflecting dynamic regulation of tumor cell migration and invasion strategies. Rap1 and Rho GTPase circuits orchestrate transitions between mesenchymal and amoeboid motility modes, which are crucial for overcoming extracellular matrix barriers during metastasis [64, 65]. Spatial scatterplots further supported these findings: within overlapping tumor regions, RAC1 (Rap1 signaling pathway) and RHOA (Cytokine-cytokine receptor interaction pathway) gene expression levels displayed a mutually inhibitory correlation pattern (Spearman’s *ρ* = -0.37), consistent with previously described models of plasticity in tumor invasion modes [66, 67].

Together, these analyses demonstrate that spatial pathway crosstalk, as revealed by Cross-Moran*^′^*s I, captures the dynamic interplay of proliferative, inflammatory, and migratory programs across tumor progression stages. By resolving not only domain-specific pathway activation but also their spatial interconnectivity, PathCLAST provides a more comprehensive view of tissue remodeling in cancer evolution.

### 3.6 Ablation study

To investigate the contributions of individual components within PathCLAST, we conducted ablation studies on the IDC dataset.

First, we evaluated the impact of the graph convolution model choice by replacing GraphSAGE with GCN. As shown in Supplementary Figure S14, this replacement led to a 17.3% decrease in clustering performance, with the ARI dropping from 0.52 to 0.43. This result indicates that the inductive representation capabilities of GraphSAGE substantially benefit the model.

Next, we systematically removed key modules from the PathCLAST architecture to assess their individual contributions (Supplementary Figure S15). The ARI values decreased as follows upon removal:

- Attention Layer-w/o: decreased to 0.38 (26.9% decrease),
- Pathway Graph-w/o: decreased to 0.34 (34.6% decrease),
- DenseNet121-w/o: decreased to 0.26 (50.0% decrease),
- Histopathological Image Inputs-w/o: decreased to 0.17 (67.3% decrease).

These findings demonstrate that each component significantly enhances the performance of PathCLAST, with the Attention Layer and Pathway Graph contributing particularly crucial roles.

We further explored optimal hyperparameter settings for contrastive learning losses and graph augmentation strategies (Supplementary Figure S16). The best performance was achieved when setting the contrastive loss weights to 0.3 for graph-to-graph (*L*_*g*2*g*_) and 0.2 for image-to-image (*L*_*i*2*i*_) losses. Regarding graph augmentation, optimal clustering was observed at a node dropping rate of 0.1 and an edge perturbation rate of 0.1.

Together, these ablation studies confirm that both the architectural design and the contrastive learning strategies are critical for achieving high performance with PathCLAST.

## 4 Discussions

PathCLAST achieved the highest clustering performance with an average ARI of 0.46 across seven different sections of the Her2ST dataset. Notably, the number of labels within each section influenced clustering performance. Sections D1, E1, and F1, which contain 3 labels each, exhibited higher ARI values compared to sections A1, B1, and G2, which contain 4, 5, and 6 labels, respectively. This suggests a weak correlation between performance and the number of labels. In section G2, label bias significantly impacted clustering performance. Sparse annotation of certain labels led to reduced clustering accuracy, as observed in sections A1, G2, where underrepresented labels were challenging to cluster accurately. We quantified the spatial distribution of labels by measuring the variance in coordinate information. A negative correlation was observed between the variance and clustering performance, indicating that greater spatial variance reduces clustering effectiveness. This effect was particularly pronounced in sections with more severe label biases (Supplementary Table S2).

The spatial intra-domain heterogeneity captured by PathCLAST offers important clinical insights into tumor progression and therapeutic resistance. While conventional histological analysis assumes relative uniformity within pathologically defined regions, our findings reveal that invasive tumors exhibit extensive spatial variation in metabolic, signaling, and immune-related pathways at sub-domain resolution. Specifically, within the invasive sub-domain (I1), the Phenylalanine metabolism and Ferroptosis pathways show both high coefficients of variation and high Moran*^′^*s I (Supplementary Figure S18). This pattern supports the known biological relationship where Phenylalanine metabolism generates BH_4_ via GCH1, which drives COQ_10_ synthesis through 4-HBA, thereby preventing lipid peroxidation and suppressing Ferroptosis [68, 69]. The suppression of ferroptosis impairs normal apoptotic cell death mechanisms, consequently promoting rapid proliferation and invasion in triple-negative breast cancer cells [70, 71]. Furthermore, in the DCIS/LCIS sub-domain (D2), both the Rap1 signaling pathway and the Focal adhesion pathway exhibit high CV and high Moran*^′^*s I (Supplementary Figure S18), indicating significant spatial heterogeneity. Activation of the Rap1 signaling pathway has been shown to facilitate talin recruitment to adhesion sites, promoting integrin activation and clustering, and leading to focal adhesion formation [72]. Under mechanical tension, these integrin clusters trigger FAK phosphorylation at Y397 and subsequent Src recruitment to form the FAK/Src complex [73, 74]. This sequence of events enhances integrinECM mechanical linkages through increased contractile force and contributes to the formation of a rigid extracellular matrix, generating mechanical heterogeneity within the tumor microenvironment [75]. As a result, these mechanotransduction processes strengthen cell adhesion, and the activated FAK/Src complex regulates cytoskeletal dynamics to promote migration and invasion, thereby driving tumor metastasis [76].

This fine-grained molecular diversity likely contributes to well-documented clinical phenomena such as variable therapeutic responses, localized resistance, and metastatic seeding from aggressive sub-clones [50, 52]. In particular, the consistent metabolic heterogeneity observed across invasive compartments may reflect region-specific adaptations to microenvironmental stresses such as hypoxia, nutrient deprivation, or immune evasion, supporting emerging models of metabolic plasticity in cancer [48, 49]. Mapping these intra-domain variations could inform precision therapeutic strategies by identifying spatial niches that harbor therapy-resistant or metastasis-initiating cells. Future integration of spatial pathway attention profiles with clinical outcome datasets may further refine risk stratification and enable spatially targeted interventions, representing a major advance in the design of next-generation spatially informed cancer therapies.

## 5 Conclusion

We introduce PathCLAST, a contrastive learning framework that integrates pathway-informed gene expression representations with histopathological image features to enhance spatial domain identification in spatial transcriptomics data. By embedding gene expression within curated pathway graphs and aligning these representations with spatially matched histological features through bi-modal contrastive learning, PathCLAST enables interpretable and biologically grounded clustering. The model demonstrates improved performance on benchmark datasets and provides mechanistic insights into tissue organization through pathway-level attention scoring and spatial autocorrelation analysis. Additionally, PathCLAST reveals intra-domain molecular heterogeneity and inter-compartmental crosstalk, highlighting spatially localized biological processes relevant to tumor progression. These capabilities make PathCLAST a valuable tool for spatial systems biology, with future applications in identifying microenvironment-specific therapeutic targets and advancing precision oncology through integration with clinical outcome data.

## Acknowledgments

This research was supported by the MSIT (Ministry of Science and ICT), Korea, under the ITRC (Information Technology Research Center) support program [IITP-2025-RS-2020-II201789], and the Artificial Intelligence Convergence Innovation Human Resources Development [IITP-2025-RS-2023-00254592], supervised by the IITP (Institute for Information & Communications Technology Planning & Evaluation). This work was supported by the National Research Foundation of Korea (NRF) grant funded by the Korea government (MSIT)[RS-2025-00560523, RS-2024-00346342]. This research was also supported by the Basic Science Research Program through the National Research Foundation of Korea (NRF) funded by the Ministry of Education [RS-2024-00393148].

## Notes

### Competing Interest Statement

The authors have declared no competing interest.

