## Supplementary Materials for "PathCLAST: Pathway-Augmented Contrastive Learning with Attention for Spatial Transcriptomics"

### PathCLAST: Pathway-Augmented Contrastive Learning with an Attention Mechanism for Improved Spatial Domain Identification in Spatial Transcriptomics Data

Minho Noh[1], Sungkyung Lee[1], Sunghyun Kim[2], Sangsoo Lim[1,*]

[1] Department of Computer Science and Artificial Intelligence, Dongguk University, 30, Pildong-ro 1-gil, Jung-gu, 04620, Seoul, South Korea

[2] Division of AI Convergence, Dongguk University, 30, Pildong-ro 1-gil, Jung-gu, 04620, Seoul, South Korea

[*] Division of Software Convergence, Dongguk University, 30, Pildong-ro 1-gil, Jung-gu, 04620, Seoul, South Korea

**Keywords:** Spatial transcriptomics; self-supervised contrastive learning; Pathway; Attention mechanism

#### Supplementary Figures

**Figure S1.** A boxplot shows ARI and NMI for eight methods (PathCLAST, STAGATE, PROST, ConGI, Hist2T, SpaGCN, stLearn, SCANPY) across the seven sections of the Her2ST dataset.
**Figure S2.** Manual annotations and comparison of spatial domains identified by SCANPY, stLearn, Hist2ST, SpaGCN, ConGI, STAGATE, PROST, and PathCLAST on the IDC dataset. The right table shows the number of spots for each label.
**Figure S3.** UMAP visualization and PAGA graphs generated by SCANPY, stLearn, Hist2ST, SpaGCN, ConGI, STAGATE, PROST, and PathCLAST embeddings respectively.
**Figure S4.** Contingency tables (n = 3,789 spots) for eight methods constructed from IDC dataset, comparing clustering density by tissue type using true and predicted domains.
**Figure S5.** Manual annotations and comparison of spatial domains identified by SCANPY, stLearn, Hist2ST, SpaGCN, ConGI, STAGATE, PROST, and PathCLAST on the 7 slices of Her2st dataset.
**Figure S6.** UMAP visualization and PAGA graphs were generated using embeddings from PathCLAST and other methods in all 7 sections of the Her2st dataset.
**Figure S7.** Comparison of spatial domains by clustering assignments by SCANPY, stLearn, Hist2ST, SpaGCN, ConGI, STAGATE, PROST, and PathCLAST manual annotation in the DLPFC dataset.
**Figure S8.** UMAP visualization and PAGA graphs were generated using embeddings from PathCLAST and other methods in all 12 sections of the DLPFC dataset.
**Figure S9.** A Pathway Attention–Based Workflow for Spatial Pattern Analysis Using MCL Clustering and Moran’s I, demonstrated on the IDC dataset. The workflow proceeds as follows: (Step 1.) construct a domain-specific spatial network, (Step 2.) perform Markov Clustering (MCL) for each domain, and (Step 3.) compute Moran’s I based on pathway attention weights in each MCL cluster to derive spatial patterns.
**Figure S10.** Pairplot and Spearman's correlation for the IDC predicted domain. The presented data indicate the correlation coefficients of Moran's I for the pathway attention weights within each predicted domain.
**Figure S11.** In the IDC dataset, a scatterplot shows the top 50 pathways by Moran’s I in each predicted domain, with grey dashed lines marking the domain-specific medians; pathways are grouped into six categories – namely EIP (Environmental Information Processing), HD (Human Diseases), CP (Cellular Processes), OS (Organismal systems), MT (Metabolism), and GIP (Genetic Information Processing).
**Figure S12.** In the IDC dataset, scatterplots show the top 10 pathways by Moran’s I for six pathway categories, with grey dashed lines indicating the category-specific medians.
**Figure S13.** Left: The diagram, a pseudo-time trajectory map, shows the direction of this trajectory, moving from DCIS/LCIS through invasive and surrounding tumor tissues toward normal tissue. Right: Visualization of the trajectory inferred by PathCLAST, with arrows indicating the path from tissue locations with low pseudo-time to those with high pseudo-time.
**Figure S14.** Ablation study on PathCLAST using GNN, GCN, and GraphSAGE. Performance for different pathway graph encoders and output-dimension settings was evaluated by ARI on the IDC dataset.
**Figure S15.**Ablation study of PathCLAST, comparing the full model with variants individually removing one module (attention layer, pretrained DenseNet – 121 weights, pathway graphs, or patch-level images). Performance was evaluated by ARI and NMI on the IDC dataset.
**Figure S16.** Ablation studies on the contrastive – loss weights ($L_{g2g}$ and $L_{i2i}$) and on graph – augmentations (node dropping and edge perturbation) were conducted on the IDC dataset, with performance evaluated by ARI.
**Fig. S17.** Scatterplots of Moran’s I (x-axis) versus coefficient of variation (CV, y-axis) for pathways in each sub-domain of the IDC dataset.
**Fig. S18.** Left: Scatterplot showing Moran's I (x-axis) plotted against the coefficient of variation (CV, y-axis) for pathways in the indicated sub-domain. Right: Top 10 pathways in that sub-domain, ranked by Moran’s I. Bottom: Spatial visualization of pathway attention weights, showing heterogeneous pathway activity across the sub-domain.

#### Supplementary Table

**Table S1.** Description of all ST datasets used in this study.
**Table S2.** Adjusted Rand index (ARI) performance of eight methods and the spot and cluster counts for each section of the Her2st, IDC, and DLPFC datasets.
**Table S3.** Pathway ANOVA table.
**Table S4.** Pathway CV table with pathway category.

### Supplementary Notes

##### Comparison with other spatial domain identification methods

We compared PathCLAST with the non-spatial clustering method implemented by SCANPY, and six recently developed spatial clustering approaches including stLearn, Hist2ST, SpaGCN, STAGATE, ConGI, and PROST The parameter settings of these methods are as follows:

SCANPY: SCANPY follows the same data preprocessing strategy as STAGATE—data is log-normalized and the top 3,000 highly variable genes are selected. Next, 30 principal components are computed, and a nearest neighbor graph is built using the default settings of the *scanpy.pp.neighbors()* function. Finally, clustering assignments were obtained using mclust for the labeled dataset (e.g., DLPFC), with the resolution parameter tuned manually to ensure that the number of clusters matches the ground truth.

stLearn: We used stLearn on the DLPFC dataset following the settings suggested in the package documentation (often called a "package tutorial"). First, we ran the function *stLearn.SME.SME_normalized()* on the raw gene counts using the parameters use_data="raw" and weights="physical_distance". Then, we took the first 50 principal components from the normalized data for further clustering and visualization. In addition, we applied the same procedure to the IDC and Her2st datasets.

SpaGCN: We applied SpaGCN to the DLPFC dataset using the recommended settings from the package documentation (often called a "package tutorial"). The same recommended parameters were also used for the IDC and Her2st datasets.

Hist2ST: We applied Hist2ST to the DLPFC dataset following the package tutorial. The histopathological image patch size was set to 112 pixels, and the expression matrix was restricted to the top 785 highly variable genes. A K-nearest-neighbors (KNN) graph with $k = 4$ was built from spot coordinates to encode spatial adjacency. Hist2ST was trained for 200 epochs, and the best checkpoint was selected via five-fold cross-validation. The same procedure was repeated for the IDC and HER2ST datasets.

STAGATE: STAGATE was applied as follows. For the DLPFC and IDC datasets, the top 3000 highly variable genes were selected using the Seurat_v3 method, and the data were normalized using total count normalization (target sum = 1e4) followed by a log transformation. Next, the spatial network was computed using the *Cal_Spatial_Net()* with a radian cutoff of 150, while the default value for $\alpha$ was maintained. For the Her2st dataset, 1000 highly variable genes were selected, and the same radian cutoff of 150 was applied.

ConGI: ConGI was applied using the recommended settings from the package documentation (often called a "package tutorial"). For the DLPFC and IDC datasets, highly variable genes were selected by choosing 1000 genes with high variability, the histopathological image patch size was set to 112, and the model was trained for 250 epochs, with the augmentation and contrastive loss weights also following the package recommendations. For the Her2st dataset, the PCA method was used to select 300 genes.

PROST: For the DLPFC and IDC datasets, PROST was applied using an adjacency matrix constructed via KNN with 24 neighbors. PCA was used to select 50 components, and a Laplacian filter (default = 2) was applied. The model was trained for 500 epochs, with iterative refinement performed three times. For the Her2st dataset, PROST was configured with 4 neighbors and PCA selecting 30 components.

| Platform | Dataset | Section | Number of clusters | Number of spots |
|---|---|---|---|---|
| Spatial Transcriptomics | Human HER2-positive breast tumor (Her2ST) | A1 | 5 | 348 |
| | | B1 | 4 | 295 |
| | | D1 | 3 | 309 |
| | | E1 | 3 | 587 |
| | | F1 | 3 | 692 |
| | | G2 | 6 | 475 |
| | | H1 | 6 | 613 |
| 10x visium | Human Breast Cancer Section 1 (IDC) | - | 4 | 3,789 |
| | Human dorsolateral prefrontal cortex (DLPFC) | 151507 | 7 | 4,226 |
| | | 151508 | 7 | 4,384 |
| | | 151509 | 7 | 4,789 |
| | | 151510 | 7 | 4,634 |
| | | 151669 | 7 | 3,661 |
| | | 151670 | 5 | 3,498 |
| | | 151671 | 5 | 4,110 |
| | | 151672 | 5 | 4,015 |
| | | 151673 | 5 | 3,639 |
| | | 151674 | 7 | 3,673 |
| | | 151675 | 7 | 3,592 |
| | | 151676 | 7 | 3,460 |

**Table S1.** Description of all ST datasets used in this study.

| Dataset | Section | Number of clusters | Number of spots | Previous Tools | | | | | | | PathCLAST (Ours) |
|---|---|---|---|---|---|---|---|---|---|---|---|
| | | | | **SCANPY** | **stLearn** | **Hist2ST** | **SpaGCN** | **STAGATE** | **ConGI** | **PROST** | |
| Her2ST | A1 | 5 | 348 | 0.19 | 0.16 | 0.43 | 0.38 | 0.31 | 0.20 | 0.11 | **0.51** |
| | B1 | 4 | 295 | 0.13 | 0.26 | 0.16 | 0.23 | 0.28 | 0.28 | 0.31 | **0.41** |
| | D1 | 3 | 309 | 0.13 | 0.09 | 0.16 | 0.29 | 0.28 | 0.38 | 0.25 | **0.57** |
| | E1 | 3 | 587 | 0.11 | 0.14 | 0.08 | 0.17 | 0.16 | 0.46 | 0.01 | **0.47** |
| | F1 | 3 | 692 | 0.15 | 0.19 | 0.23 | 0.12 | 0.17 | **0.43** | 0.26 | 0.41 |
| | G2 | 6 | 475 | 0.11 | 0.24 | 0.13 | 0.18 | 0.30 | 0.23 | 0.26 | **0.32** |
| | H1 | 6 | 613 | 0.19 | 0.43 | 0.24 | 0.34 | 0.40 | 0.40 | 0.28 | **0.52** |
| IDC | - | 4 | 3,789 | 0.14 | 0.21 | 0.18 | 0.13 | 0.38 | 0.31 | 0.22 | **0.52** |
| DLPFC | 151507 | 7 | 4,226 | 0.23 | 0.21 | 0.18 | 0.30 | **0.47** | 0.40 | **0.47** | 0.40 |
| | 151508 | 7 | 4,384 | 0.10 | 0.32 | 0.25 | 0.30 | **0.49** | 0.42 | 0.45 | 0.32 |
| | 151509 | 7 | 4,789 | 0.15 | 0.26 | 0.12 | 0.21 | 0.38 | 0.35 | **0.40** | **0.40** |
| | 151510 | 7 | 4,634 | 0.09 | 0.27 | 0.17 | 0.27 | **0.39** | 0.34 | 0.36 | 0.36 |
| | 151669 | 7 | 3,661 | 0.20 | 0.17 | 0.26 | 0.27 | 0.38 | 0.34 | 0.38 | **0.40** |
| | 151670 | 5 | 3,498 | 0.15 | 0.31 | 0.12 | 0.13 | 0.37 | 0.22 | 0.36 | **0.40** |
| | 151671 | 5 | 4,110 | 0,15 | 0.30 | 0.19 | 0.27 | 0.32 | **0.48** | **0.48** | **0.48** |
| | 151672 | 5 | 4,015 | 0,17 | 0.36 | 0.16 | 0.20 | 0.44 | 0.42 | **0.49** | 0.45 |
| | 151673 | 5 | 3,639 | 0.15 | 0.33 | 0.19 | 0.28 | 0.43 | 0.34 | **0.49** | 0.35 |
| | 151674 | 7 | 3,673 | 0.18 | **0.42** | 0.27 | 0.28 | 0.38 | 0.35 | 0.35 | 0.31 |
| | 151675 | 7 | 3,592 | 0.16 | **0.40** | 0.15 | 0.30 | 0.33 | 0.35 | 0.38 | 0.34 |
| | 151676 | 7 | 3,460 | 0.19 | 0.38 | 0.25 | 0.30 | **0.45** | 0.35 | 0.39 | 0.36 |

**Table S2.** Adjusted Rand index (ARI) performance of eight methods and the spot and cluster counts for each section of the Her2st, IDC, and DLPFC datasets.

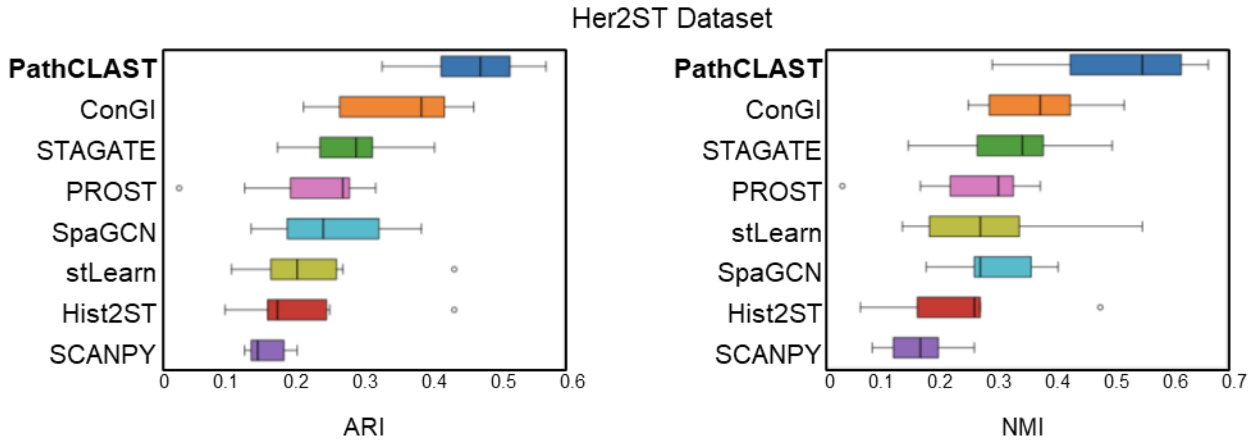


**Figure S1.** A boxplot shows ARI and NMI for eight methods (PathCLAST, STAGATE, PROST, ConGI, Hist2T, SpaGCN, stLearn, SCANPY) across the seven sections of the Her2ST dataset.

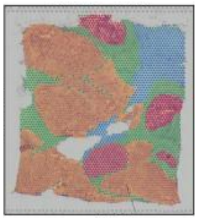


| Label | Number of Spots |
|---|---|
| Invasive | 2,027 |
| DCIS/LCIS | 463 |
| Surrounding Tumor | 823 |
| Healthy | 485 |
| Total | 3,798 |

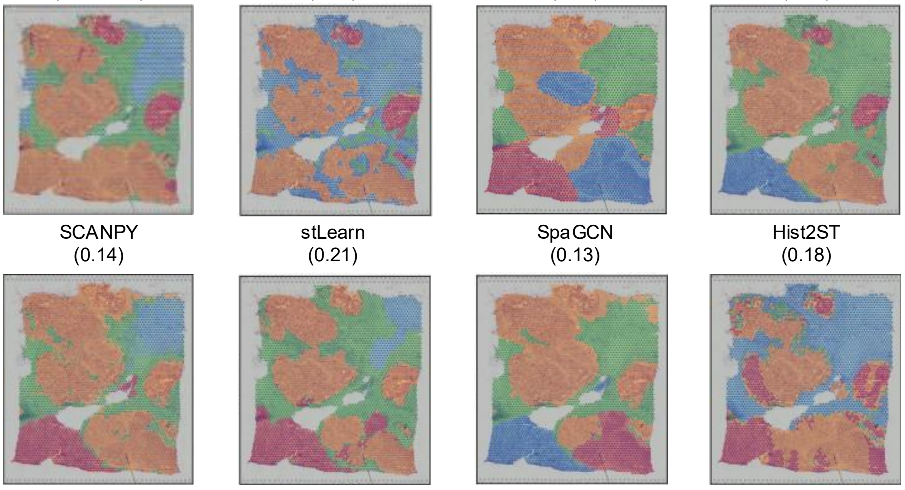


**Figure S2.** Manual annotations and comparison of spatial domains identified by SCANPY, stLearn, Hist2ST, SpaGCN, ConGI, STAGATE, PROST, and PathCLAST on the IDC dataset. The right table shows the number of spots for each label.

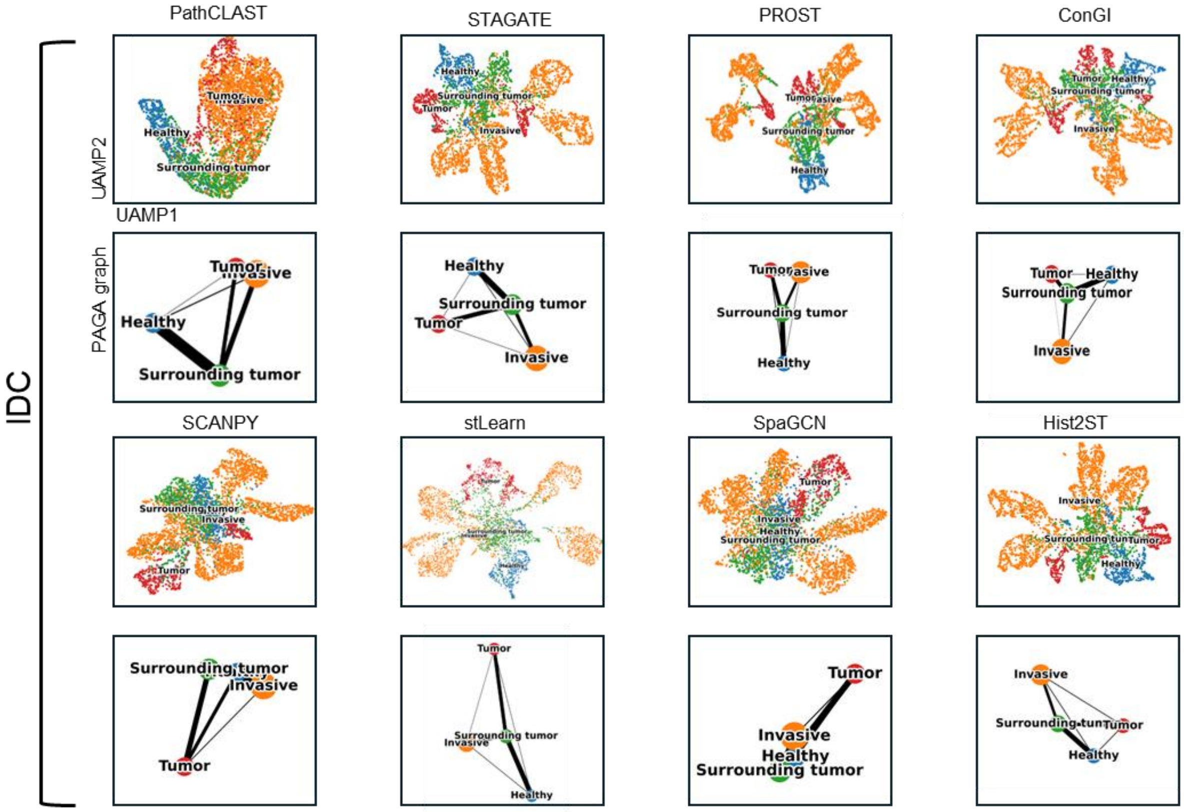


**Figure S3.** UMAP visualization and PAGA graphs generated by SCANPY, stLearn, Hist2ST, SpaGCN, ConGI, STAGATE, PROST, and PathCLAST embeddings respectively.

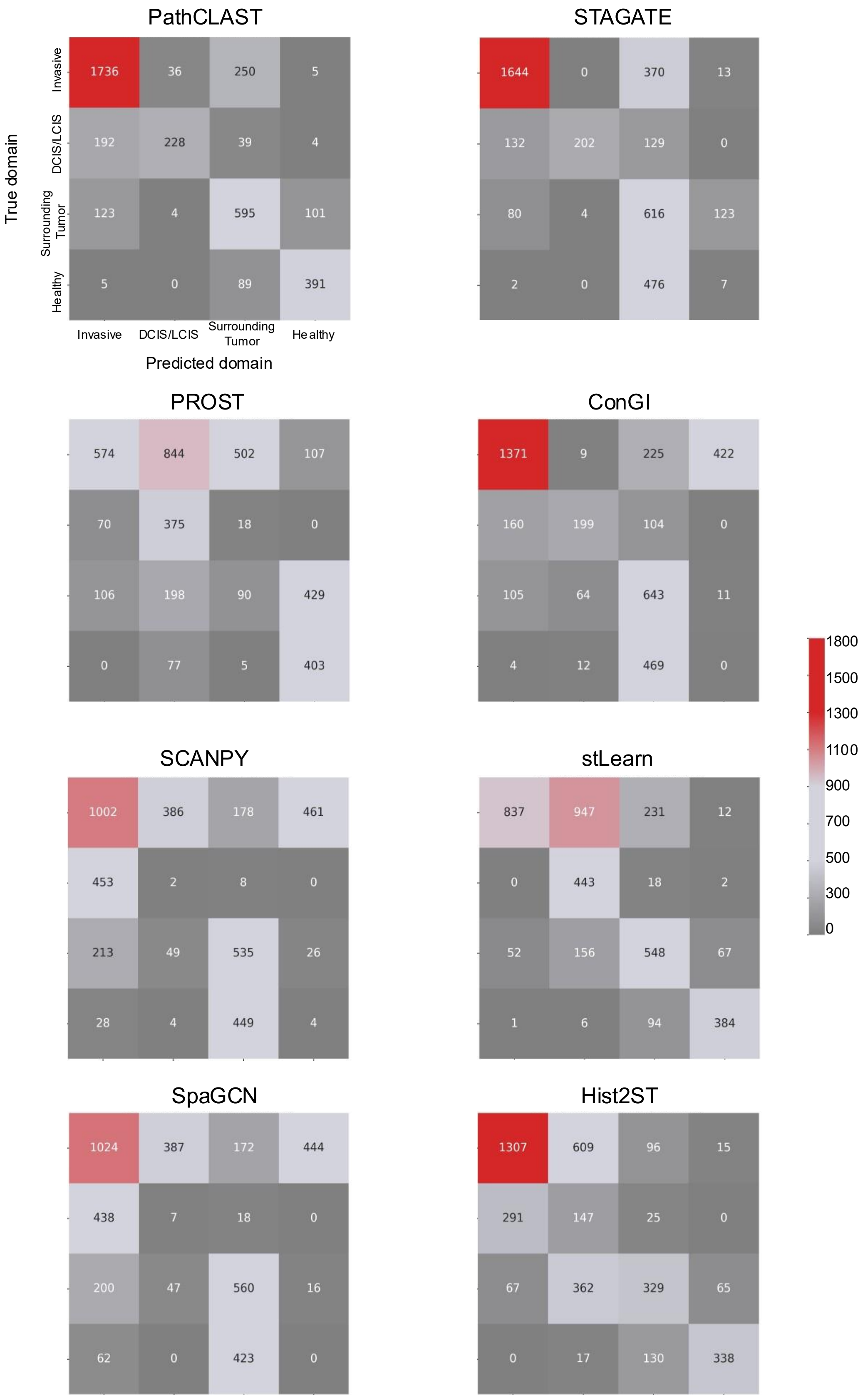


**Figure S4.** Contingency tables (n = 3,789 spots) for eight methods constructed from IDC dataset, comparing clustering density by tissue type using true and predicted domains.

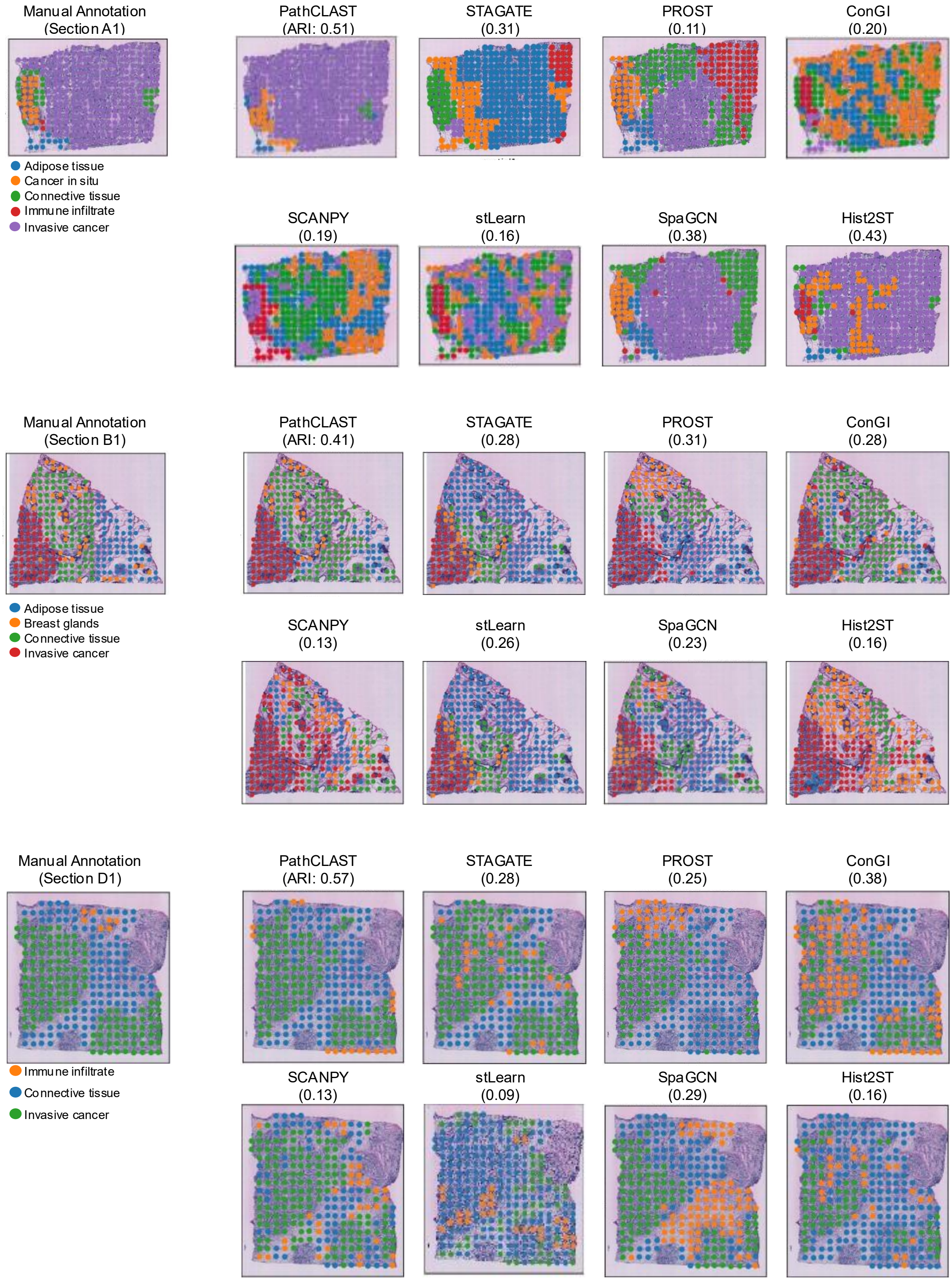


See next page

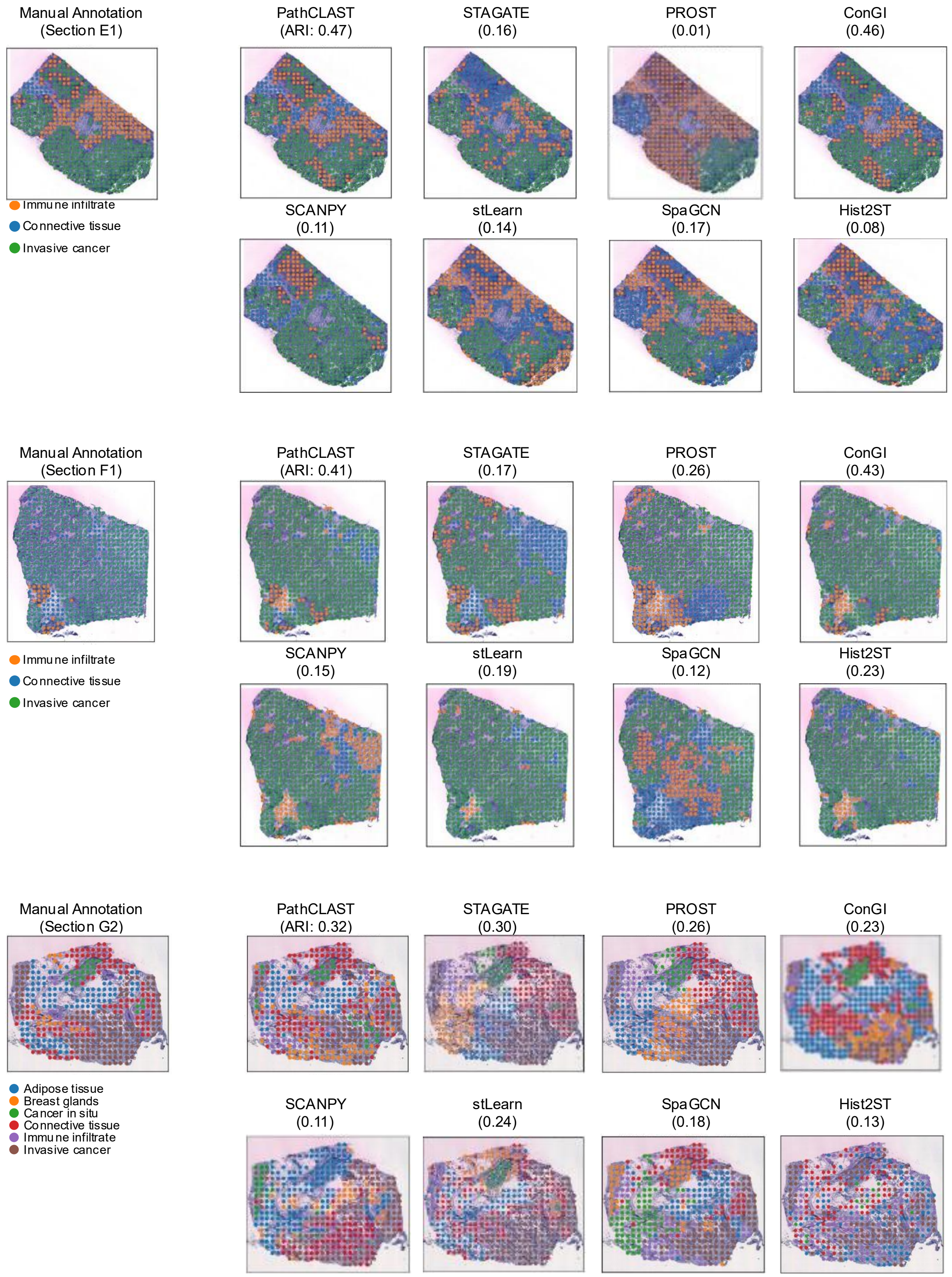


See next page

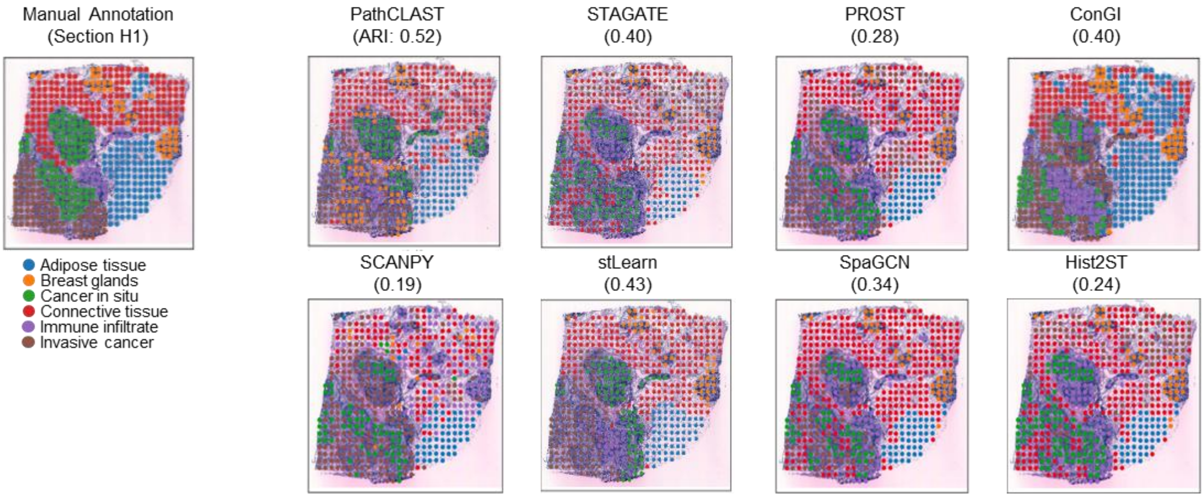


**Figure S5.** Manual annotations and comparison of spatial domains identified by SCANPY, stLearn, Hist2ST, SpaGCN, ConGI, STAGATE, PROST, and PathCLAST on the 7 sections of Her2ST dataset.

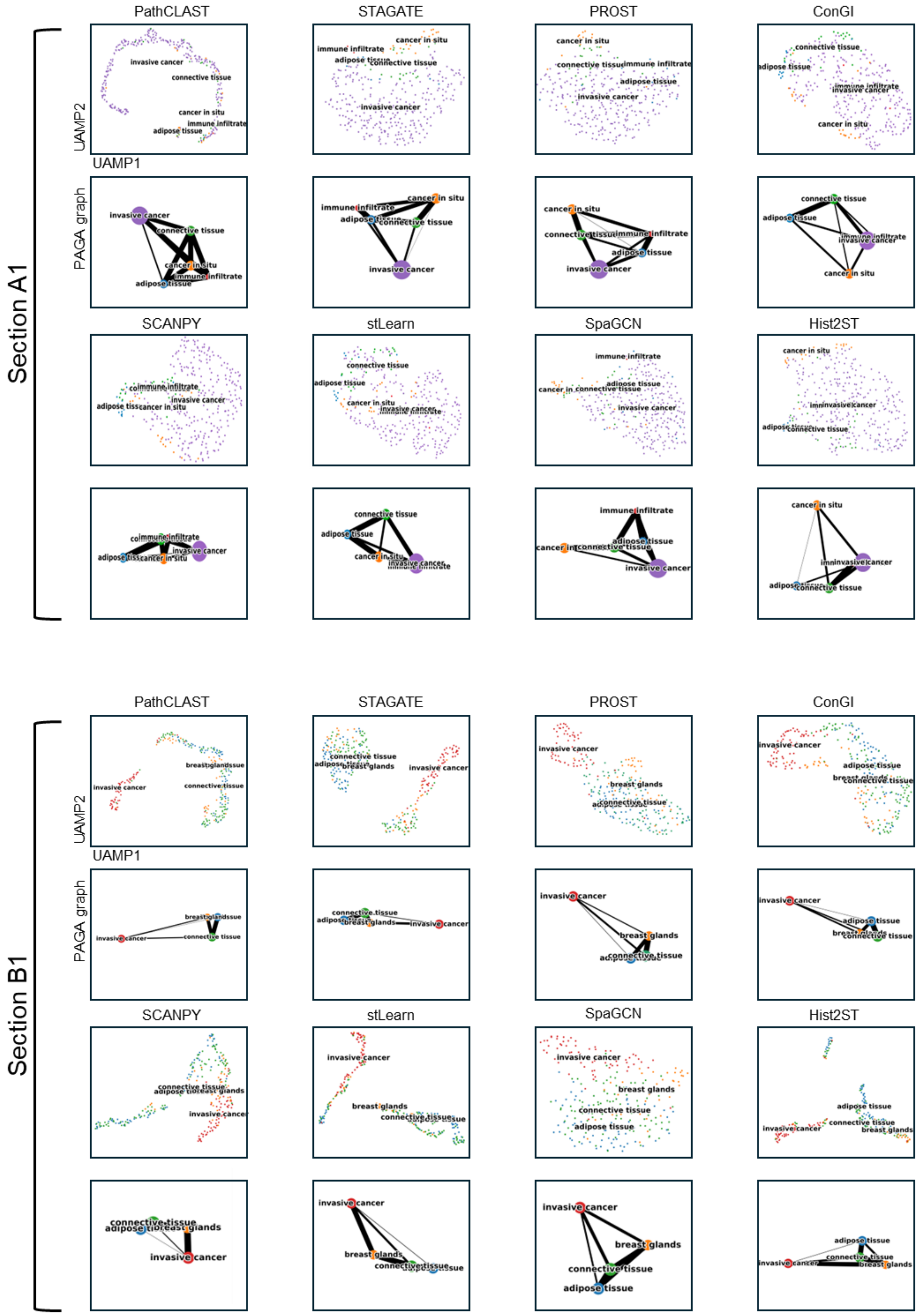


See next page

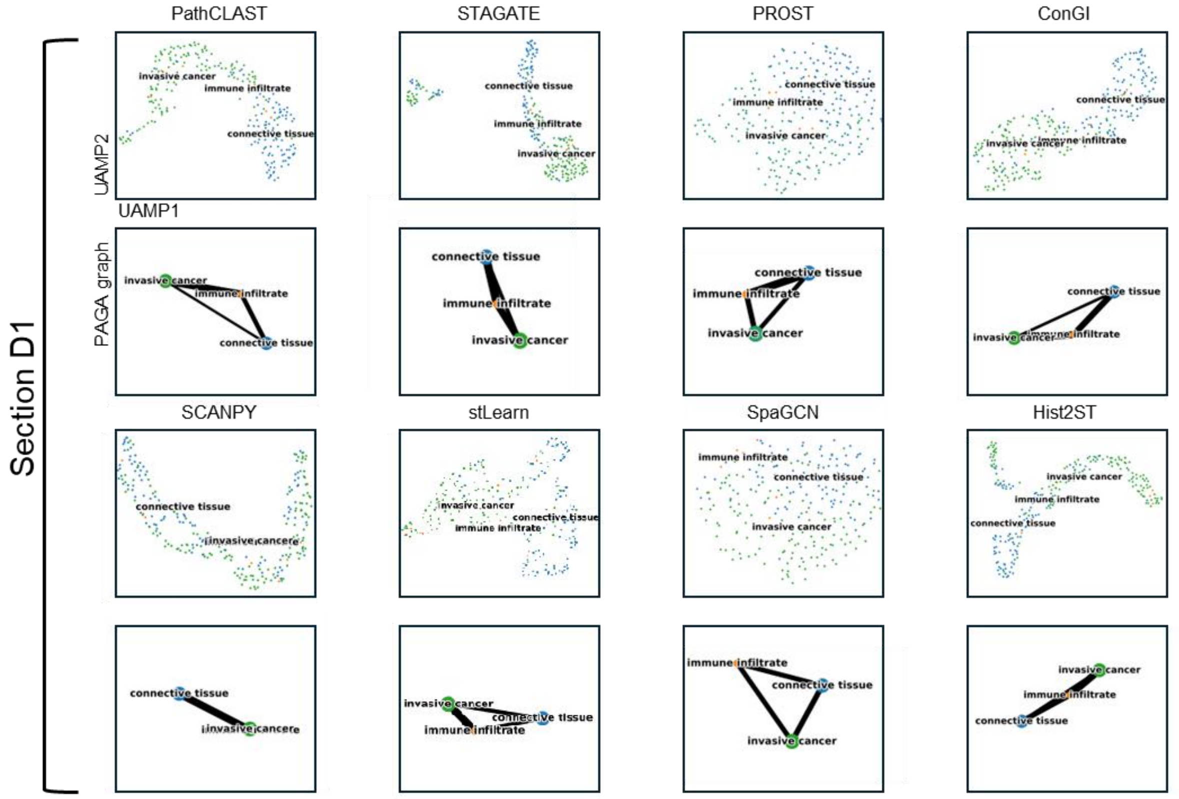


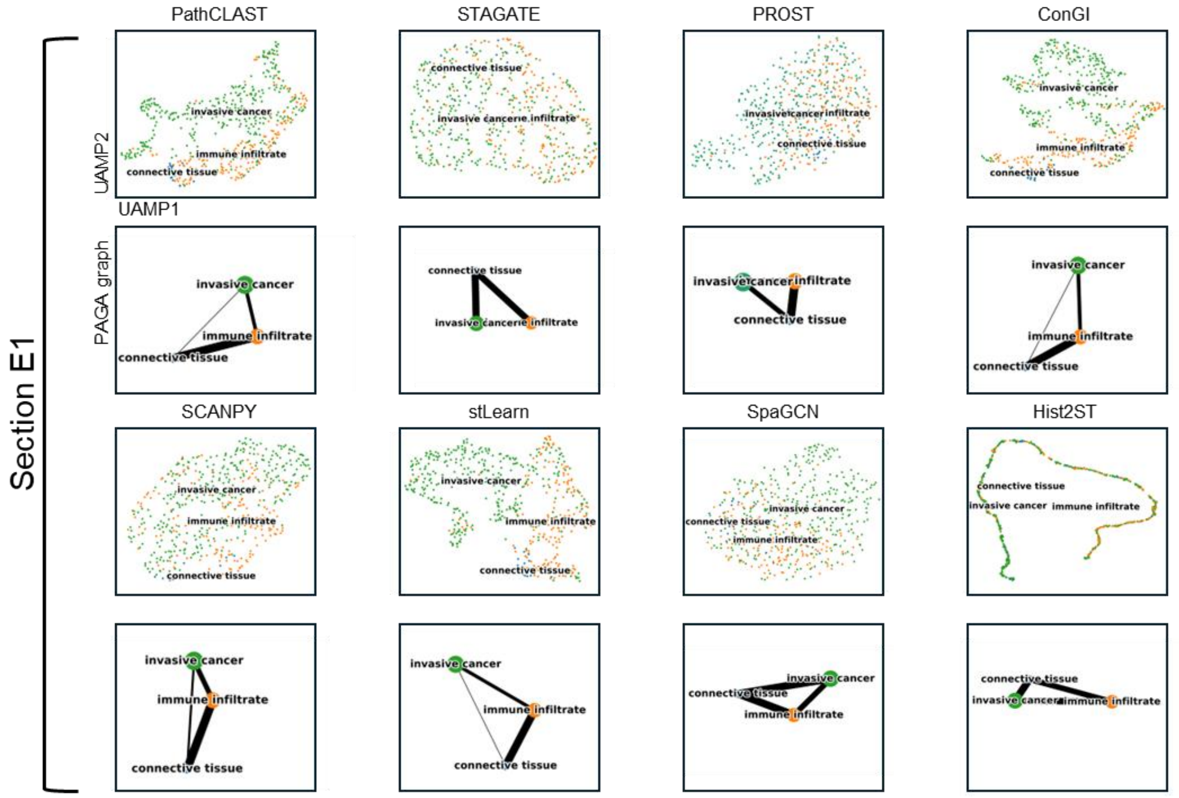


See next page

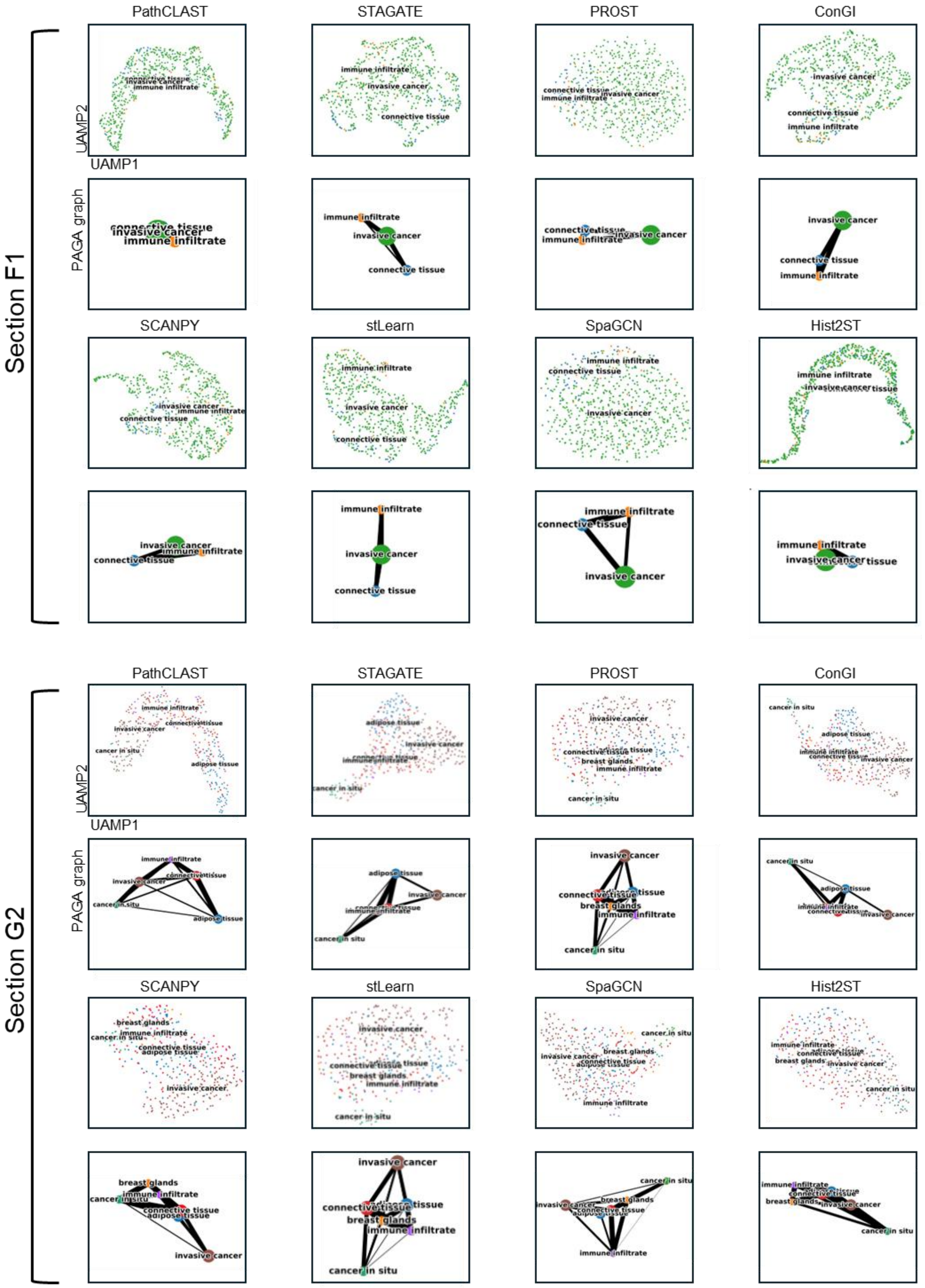


See next page

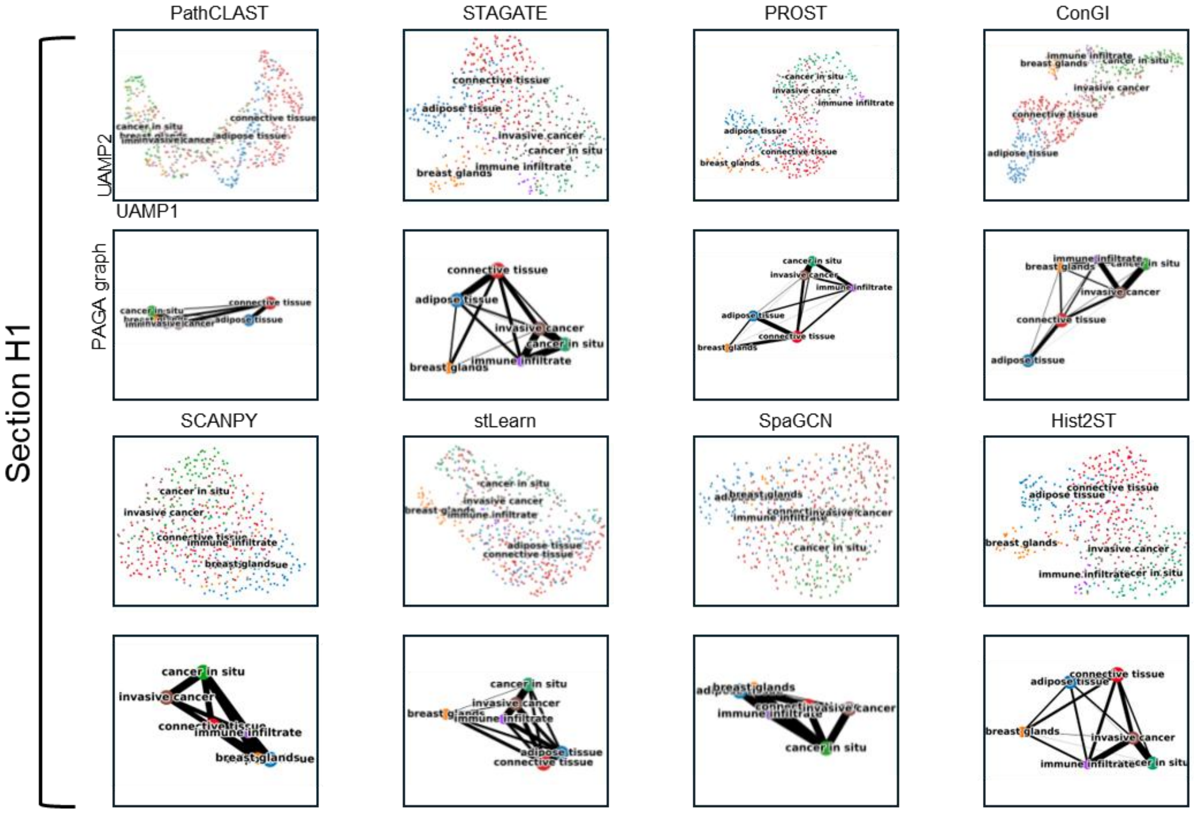


**Figure S6.** UMAP visualization and PAGA graphs were generated using embeddings from PathCLAST and other methods in all 7 sections of the Her2ST dataset.

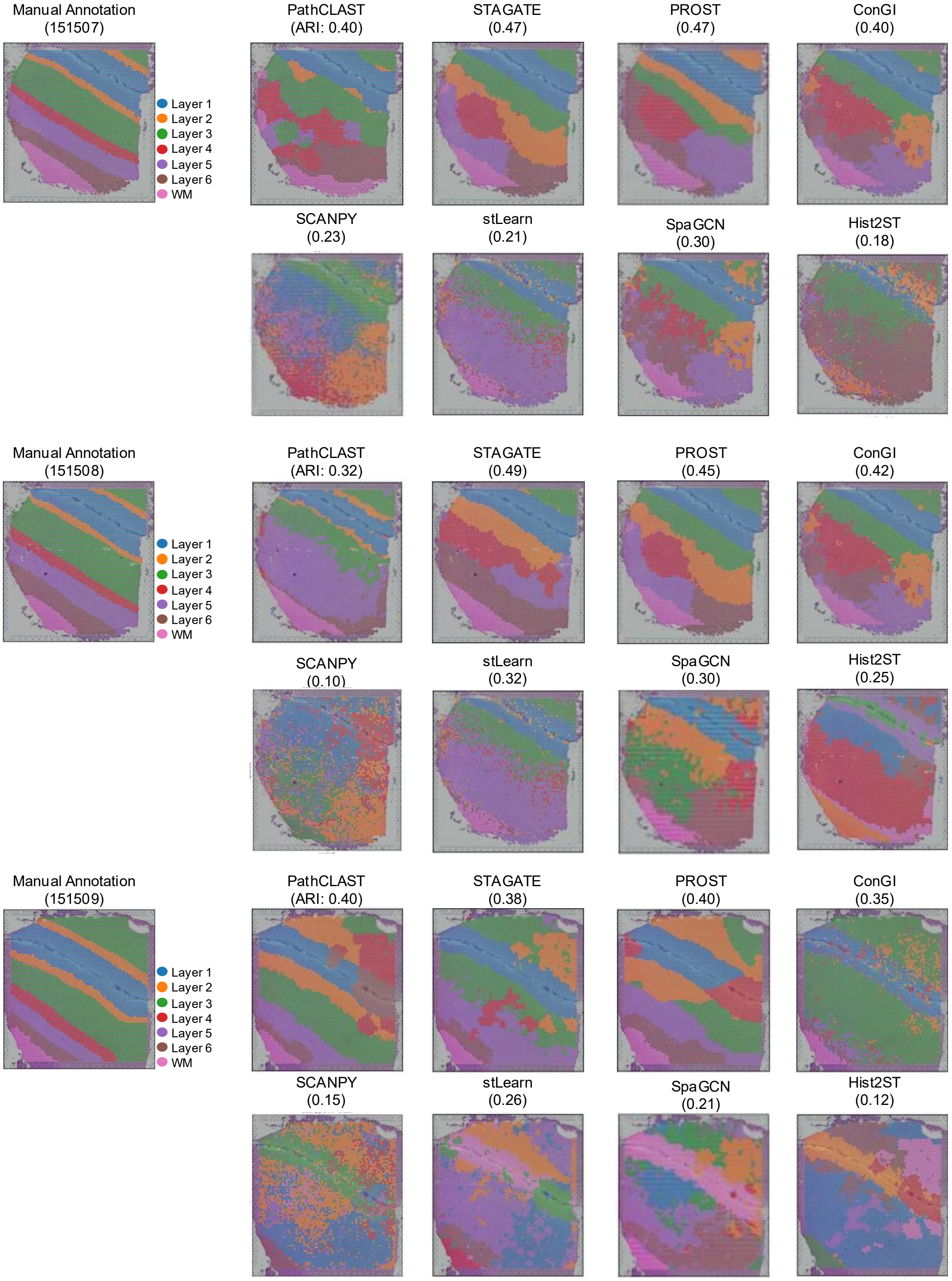


See next page

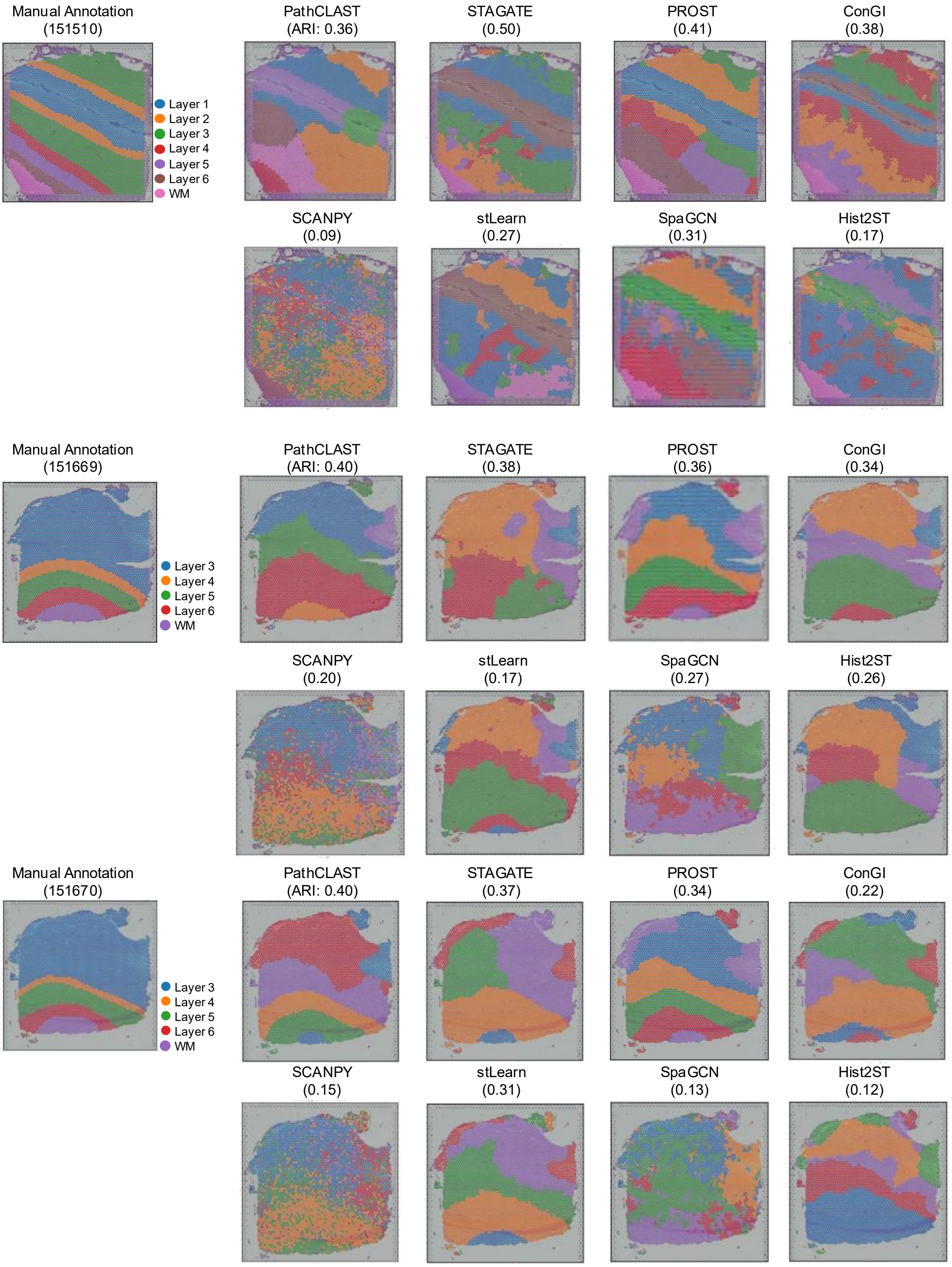


See next page

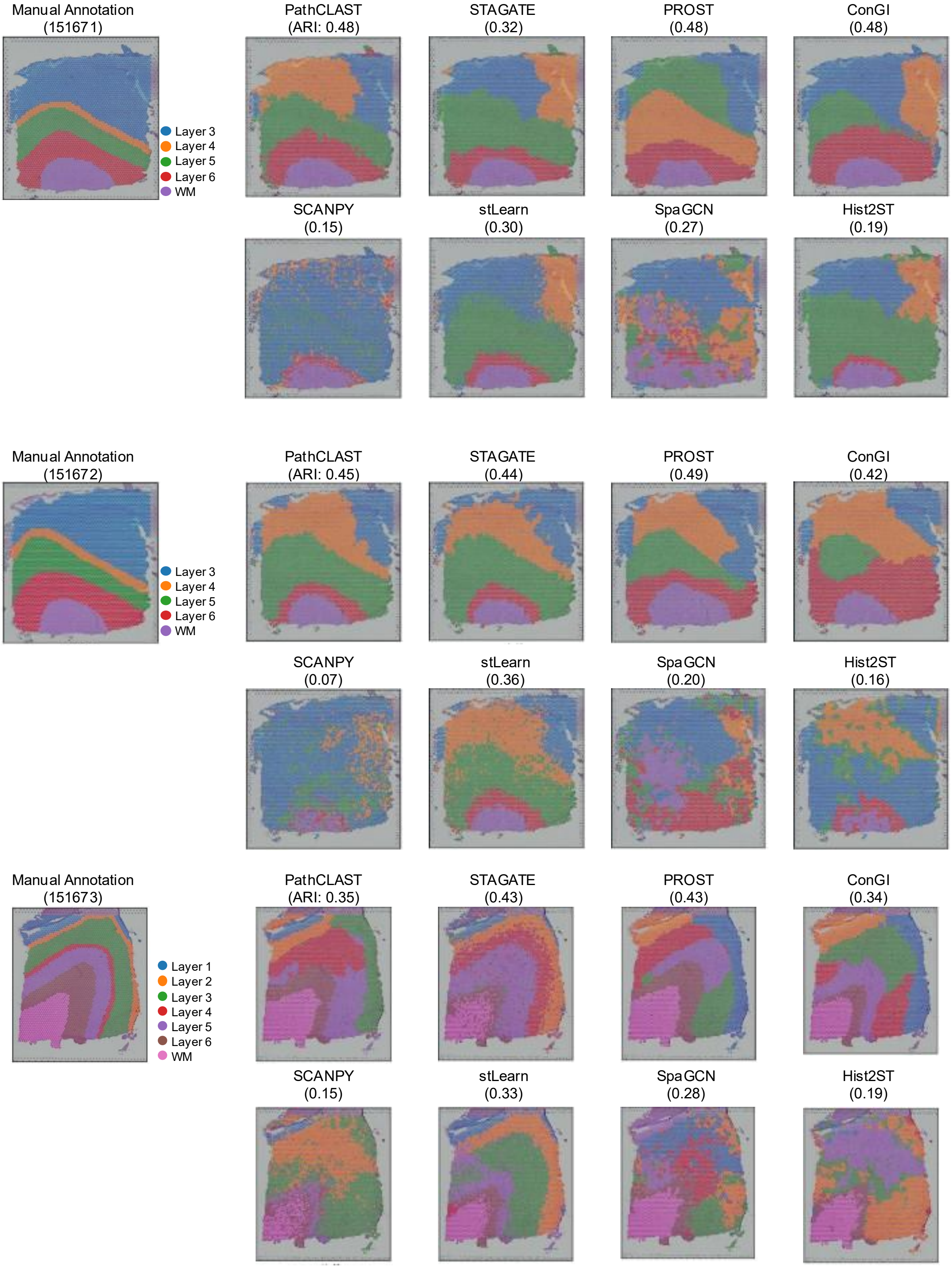


See next page

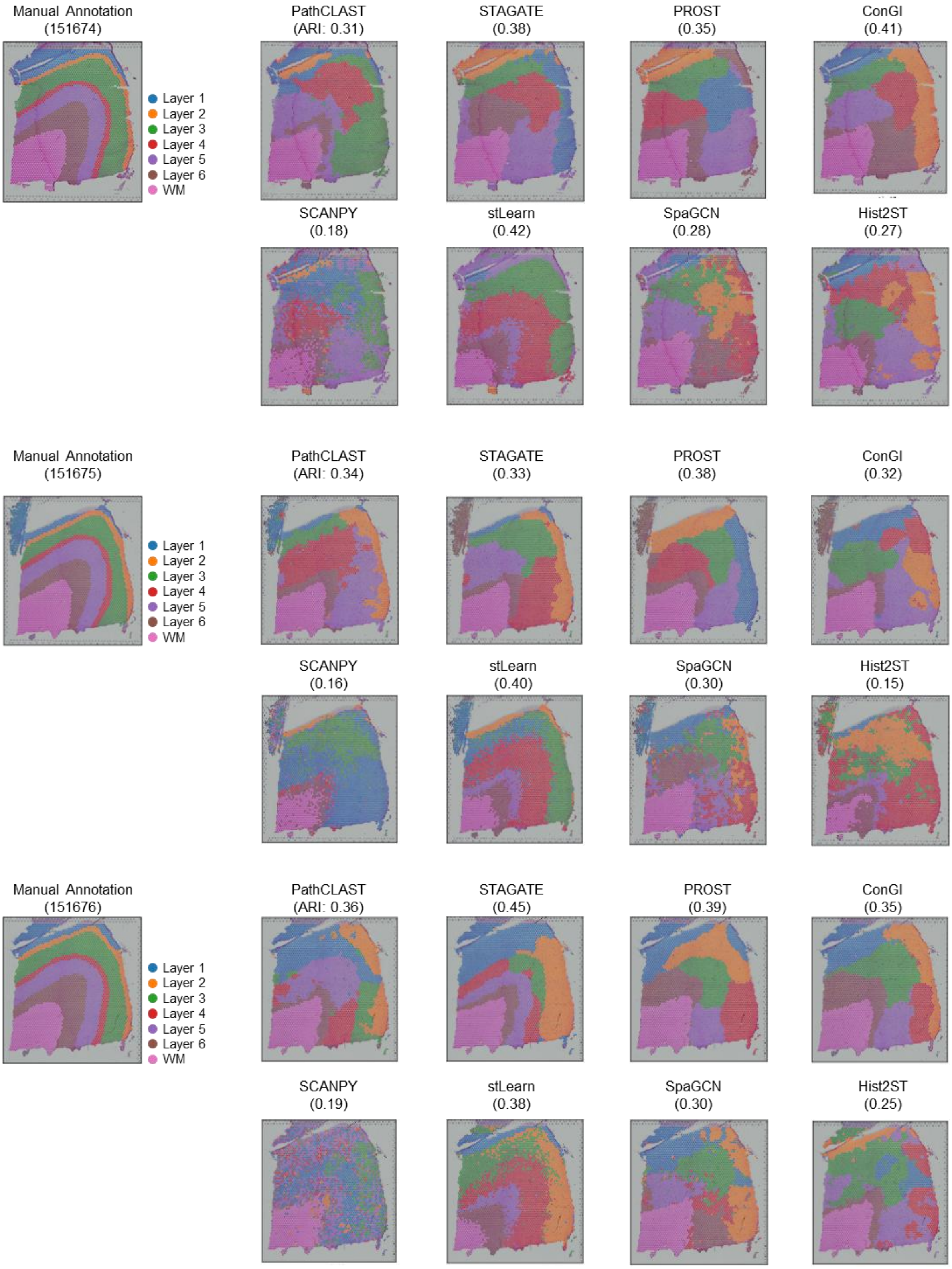


**Figure S7.** Comparison of spatial domains by clustering assignments using PathCLAST, other methods, and manual annotation in all 12 sections of the DLPFC dataset.

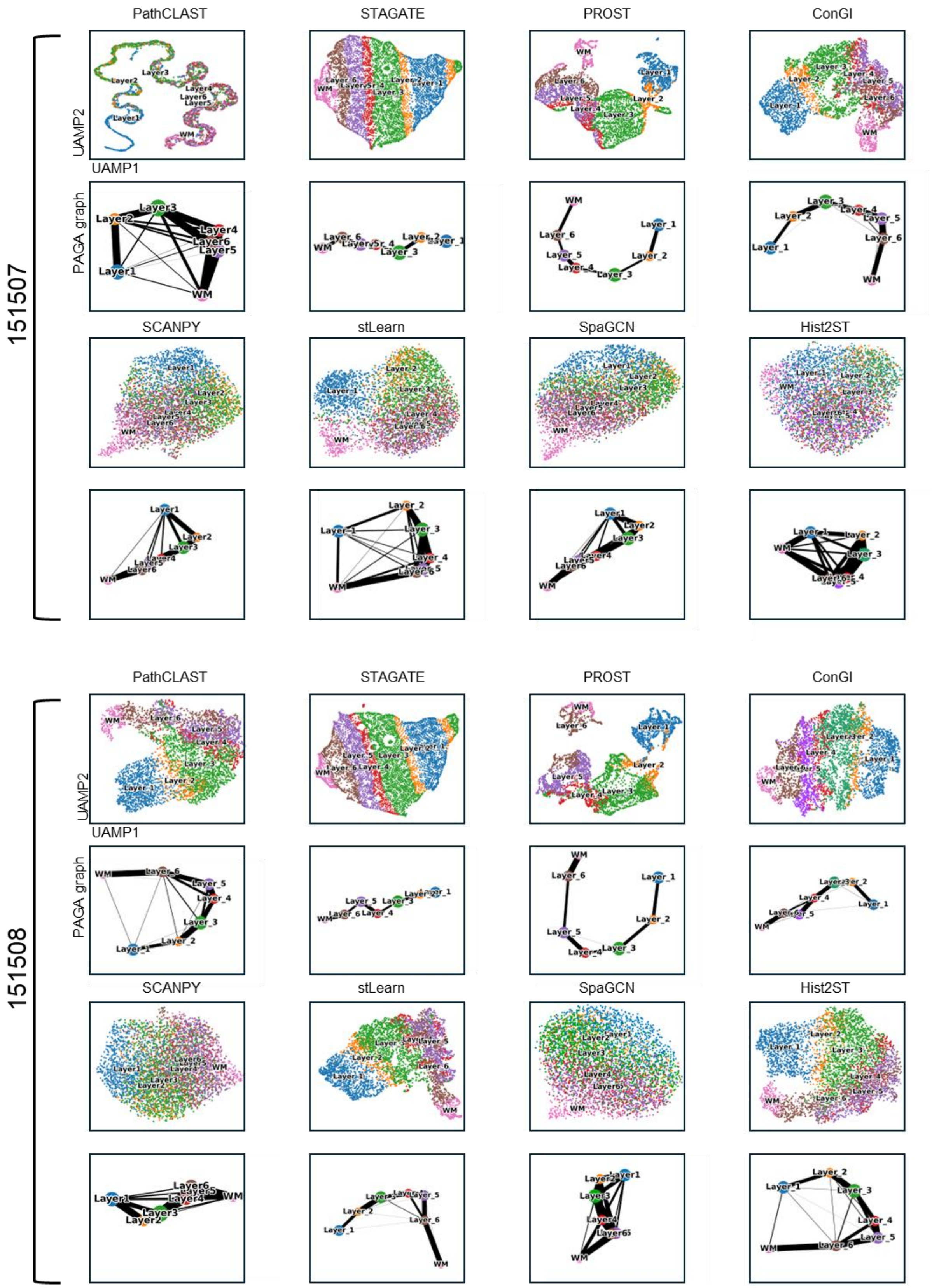


See next page

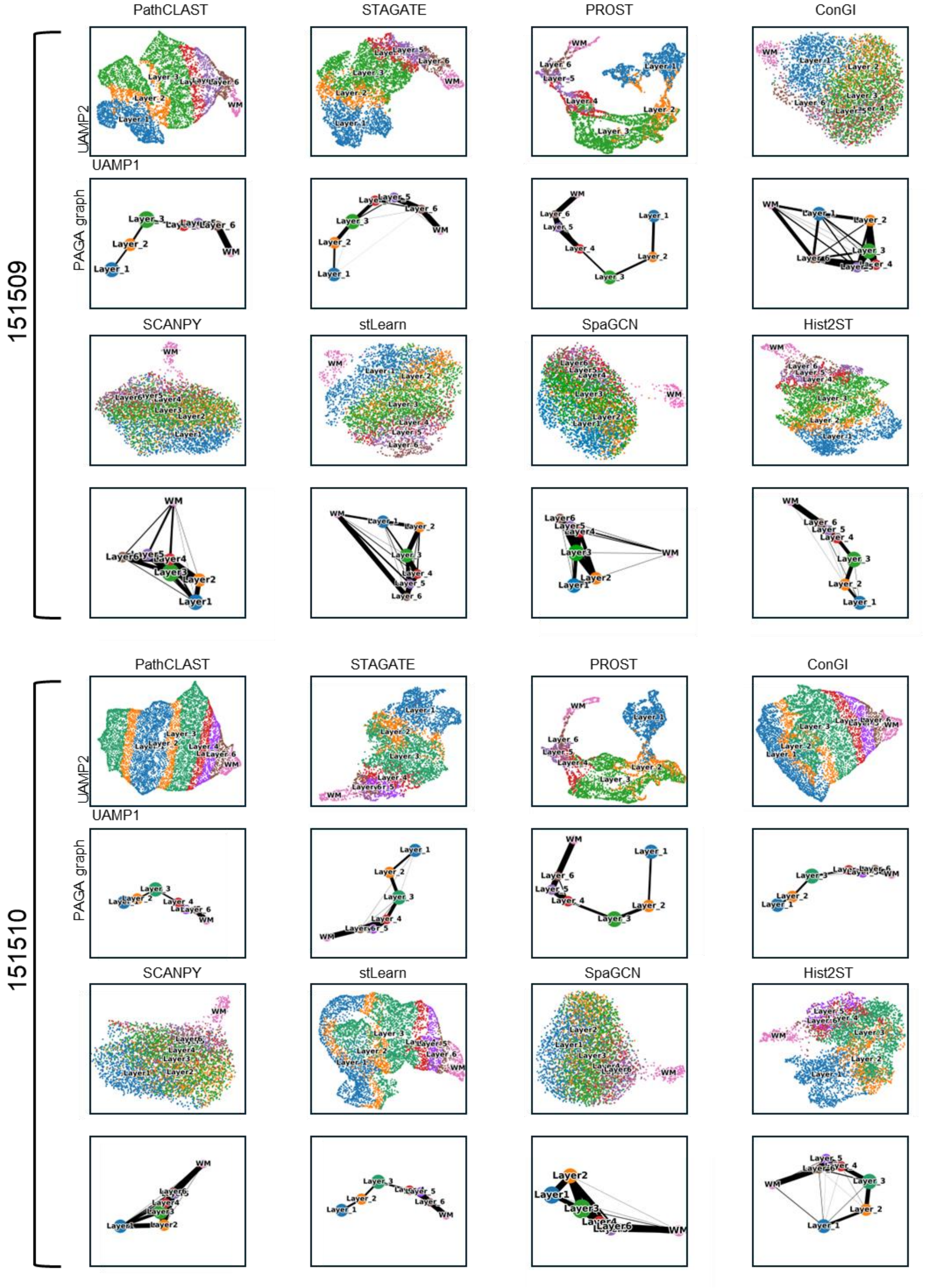


See next page

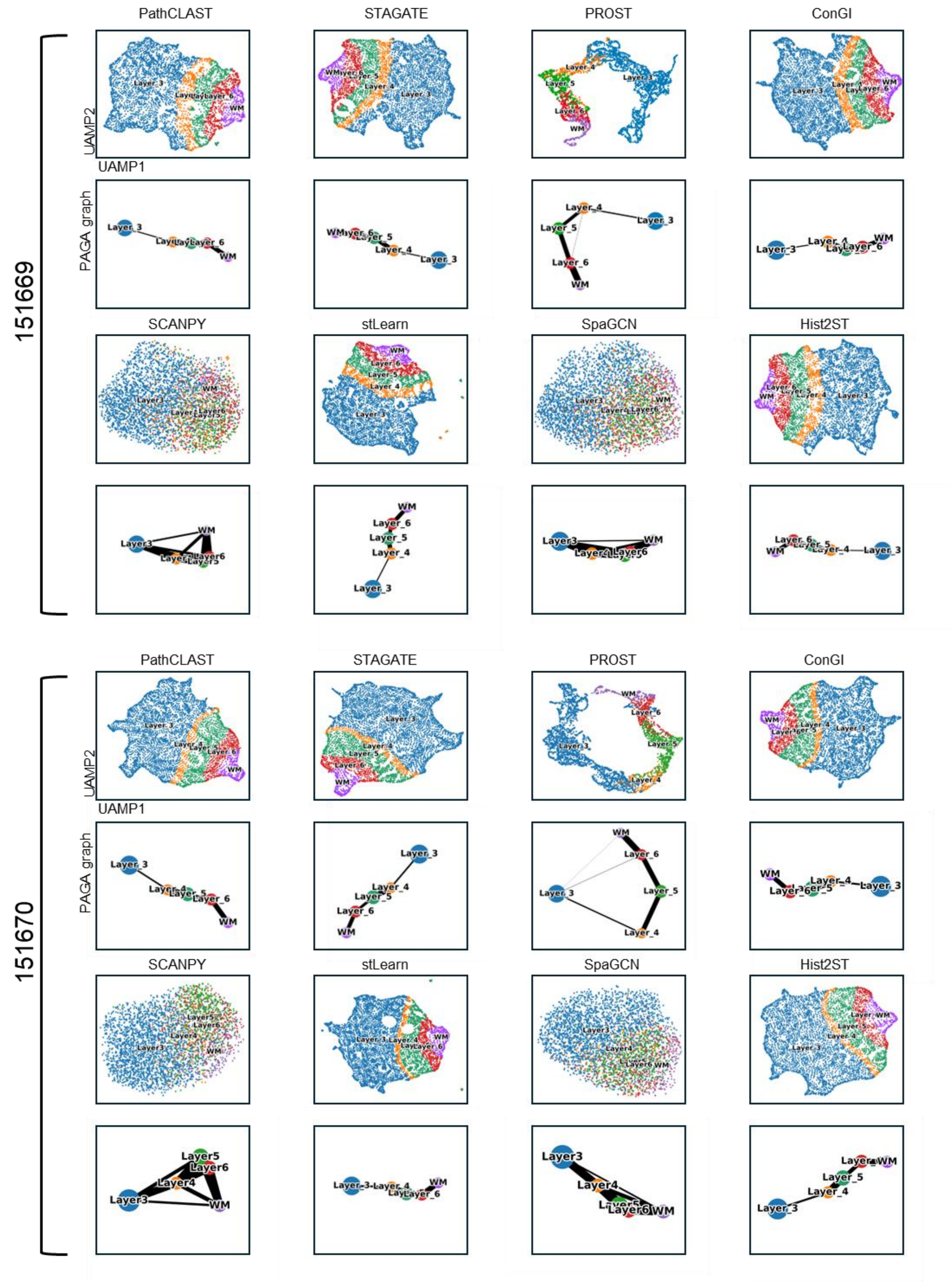


See next page

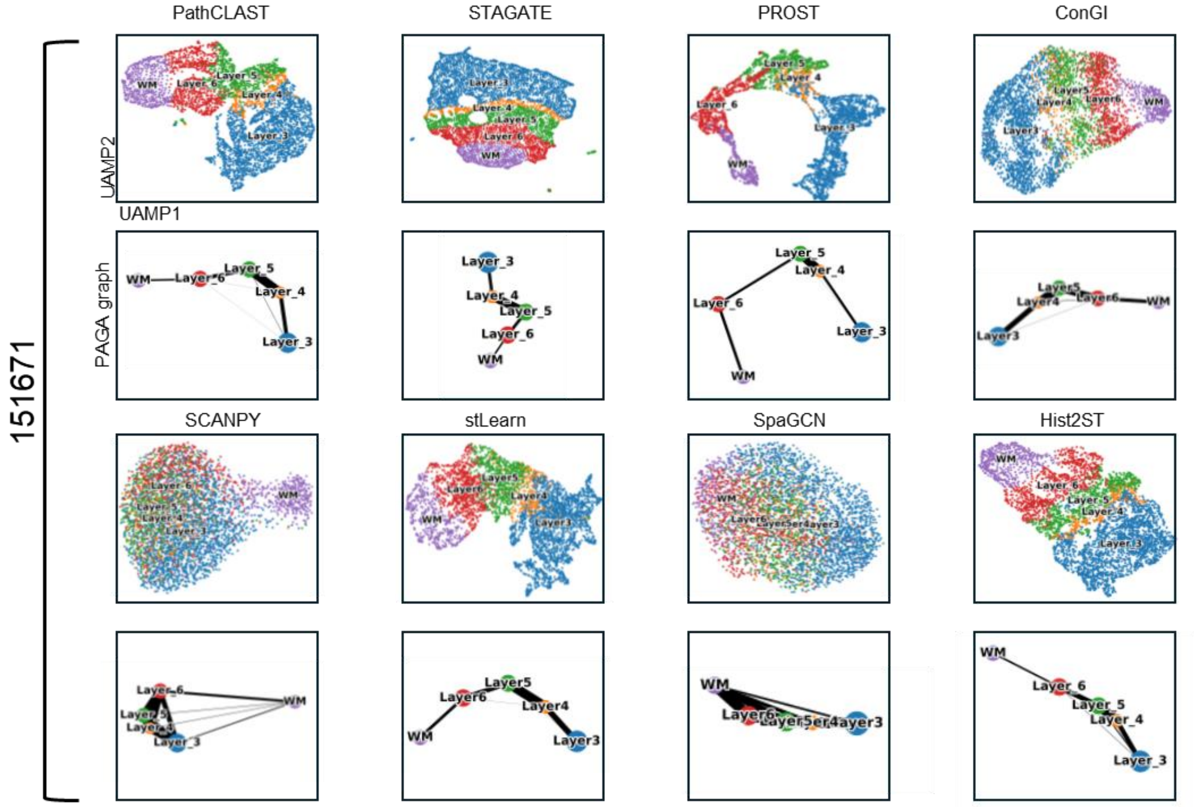


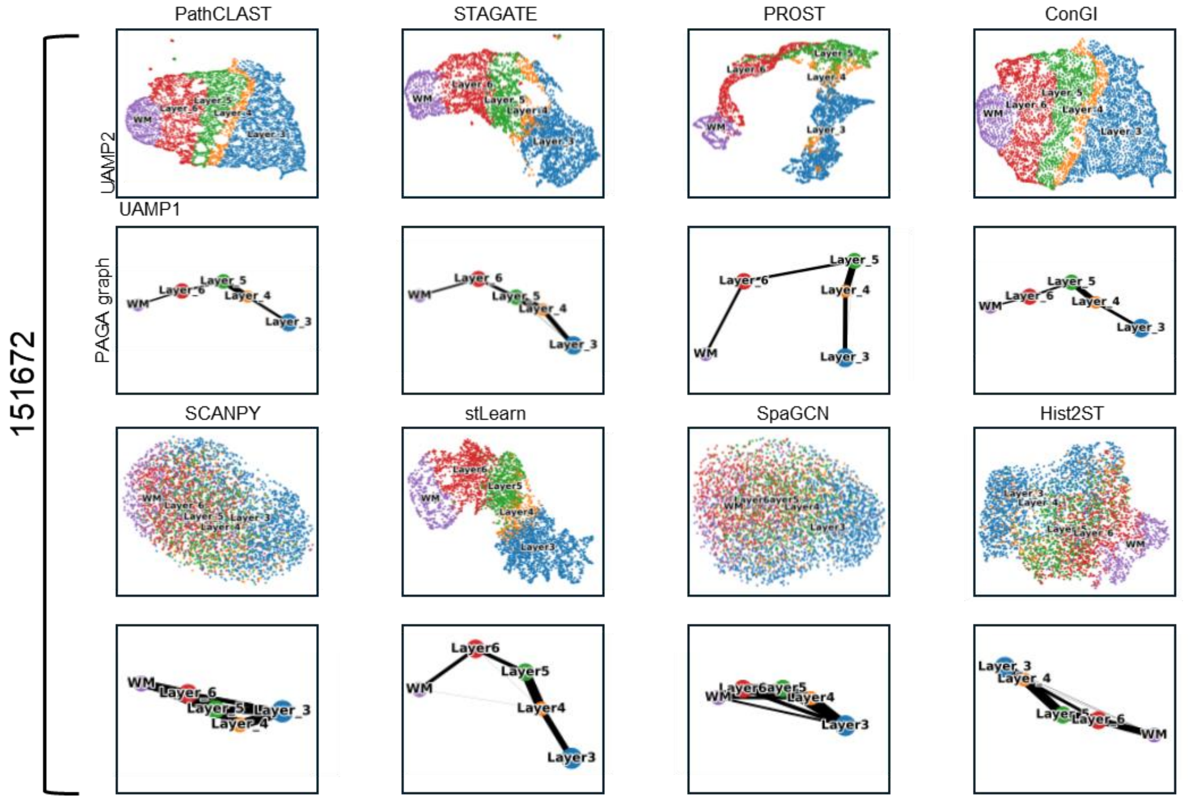


See next page

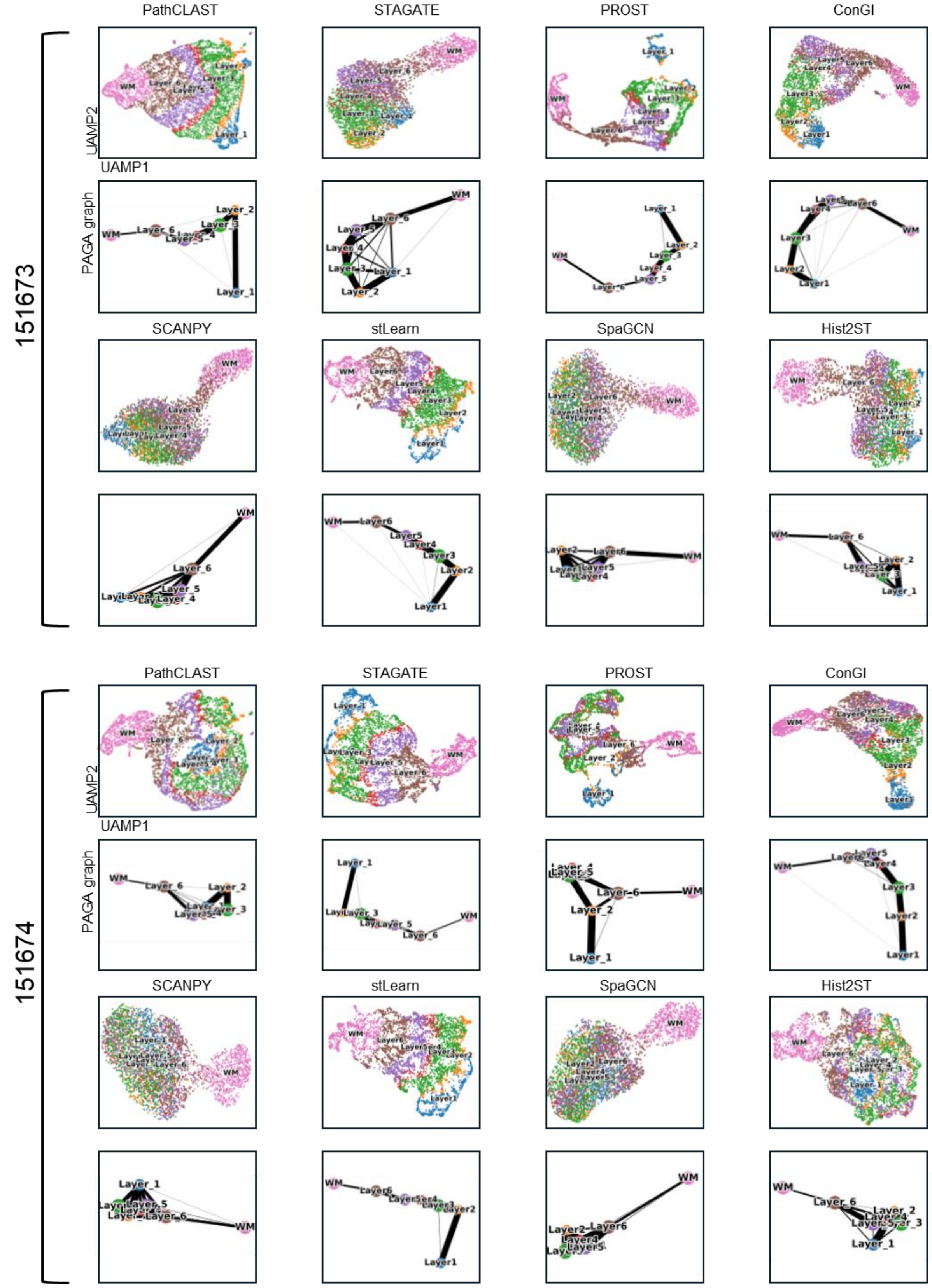


See next page

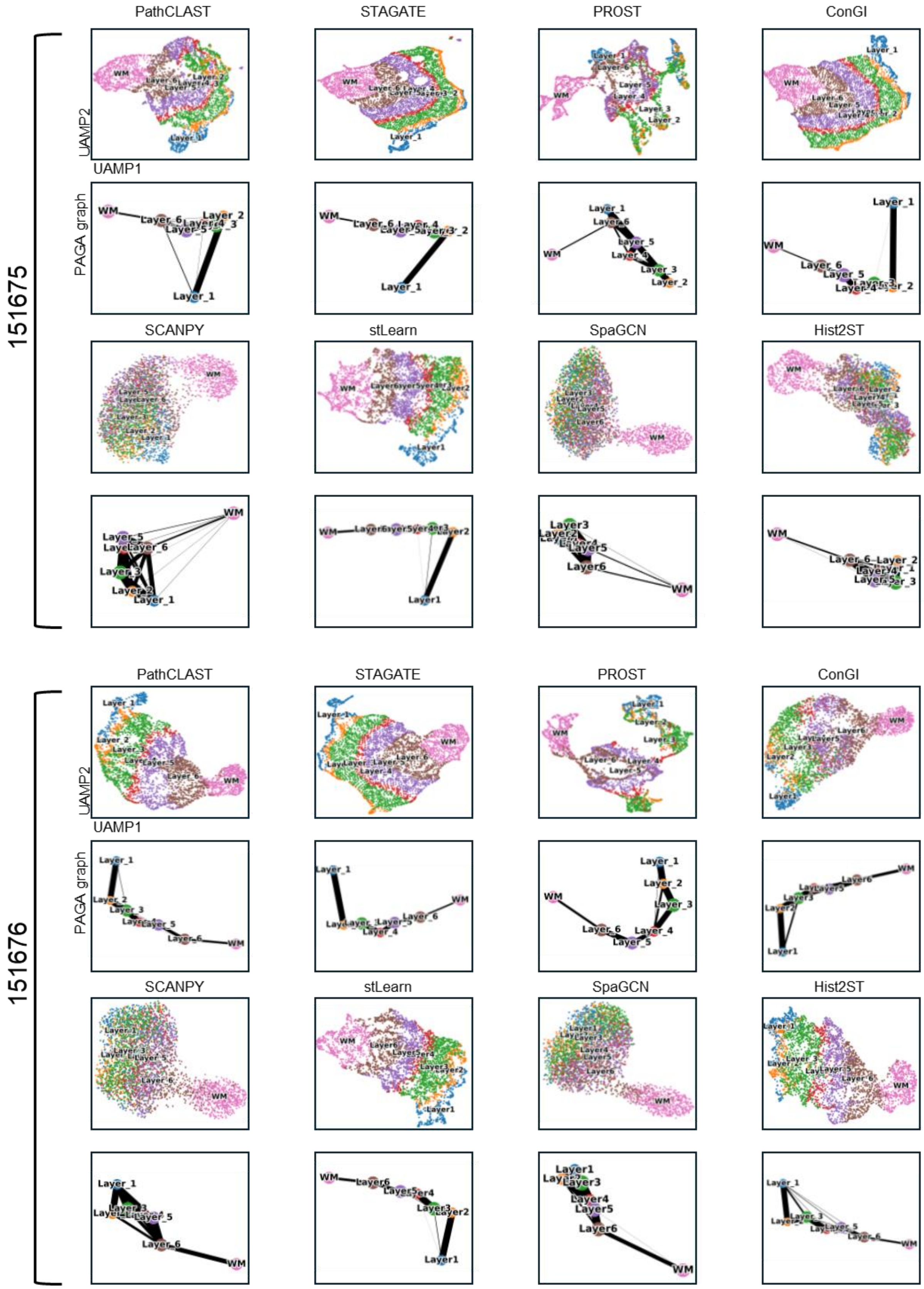


**Figure S8.** UMAP visualization and PAGA graphs were generated using embeddings from PathCLAST and other methods in all 12 sections of the DLPFC dataset.

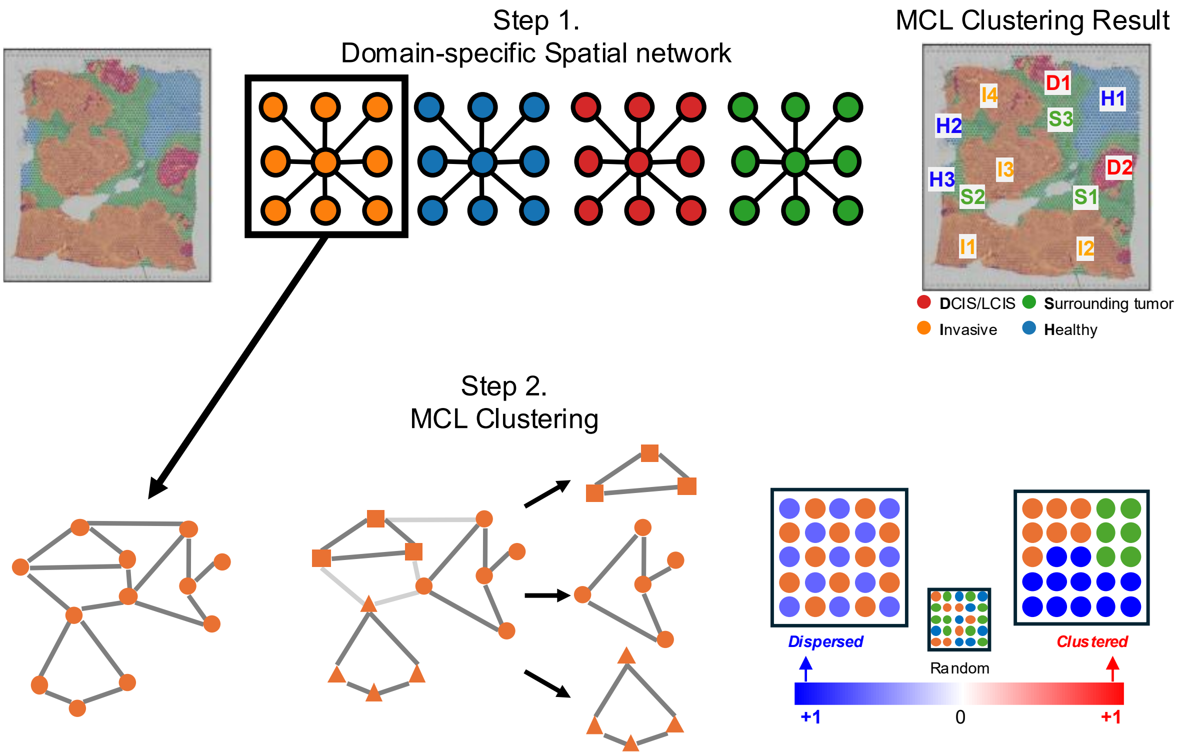


**Figure S9.** A Pathway Attention–Based Workflow for Spatial Pattern Analysis Using MCL Clustering and Moran's I, demonstrated on the IDC dataset. The workflow proceeds as follows: (Step 1.) construct a domain-specific spatial network, (Step 2.) perform Markov Clustering (MCL) for each domain, and (Step 3.) compute Moran's I based on pathway attention weights in each MCL cluster to derive spatial patterns.

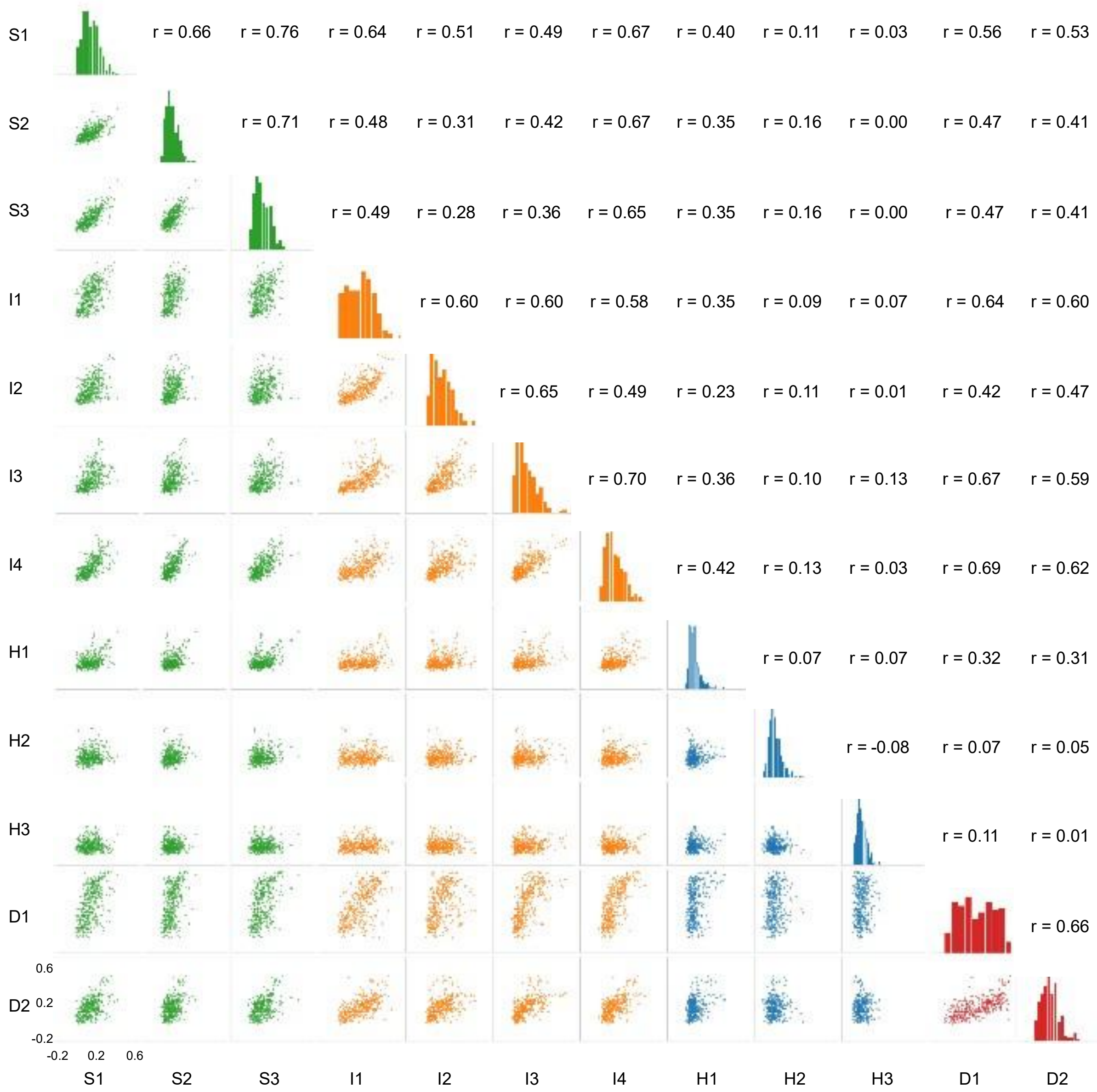


**Figure S10.** Pairplot and Spearman's correlation for the IDC predicted domain. The presented data indicate the correlation coefficients of Moran's I for the pathway attention weights within each predicted domain.

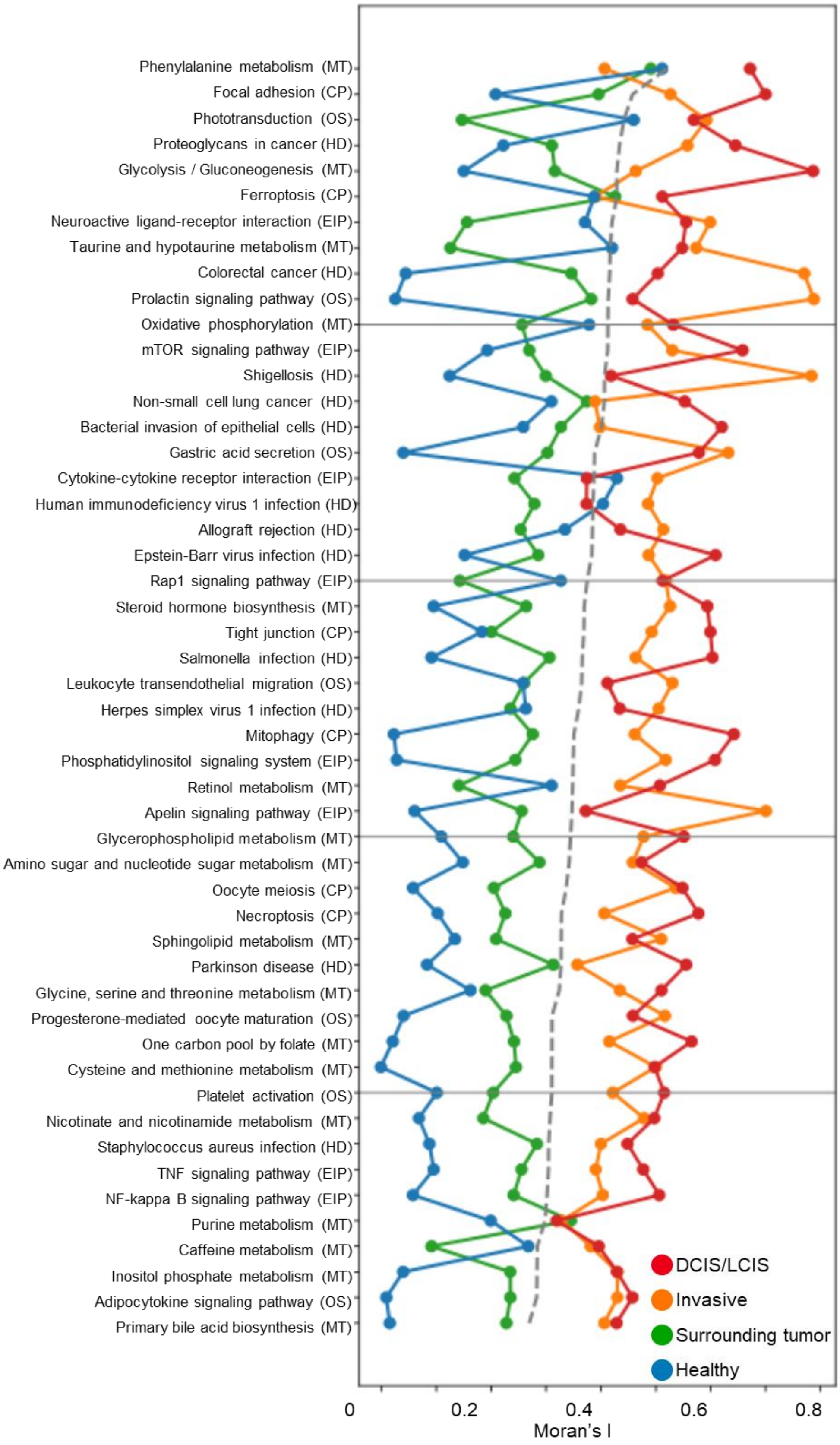


**Figure S11.** In the IDC dataset, a scatterplot shows the top 50 pathways by Moran's I in each predicted domain, with grey dashed lines marking the domain-specific medians; pathways are grouped into six categories – namely EIP (Environmental Information Processing), HD (Human Diseases), CP (Cellular Processes), OS (Organismal systems), MT (Metabolism), and GIP (Genetic Information Processing).

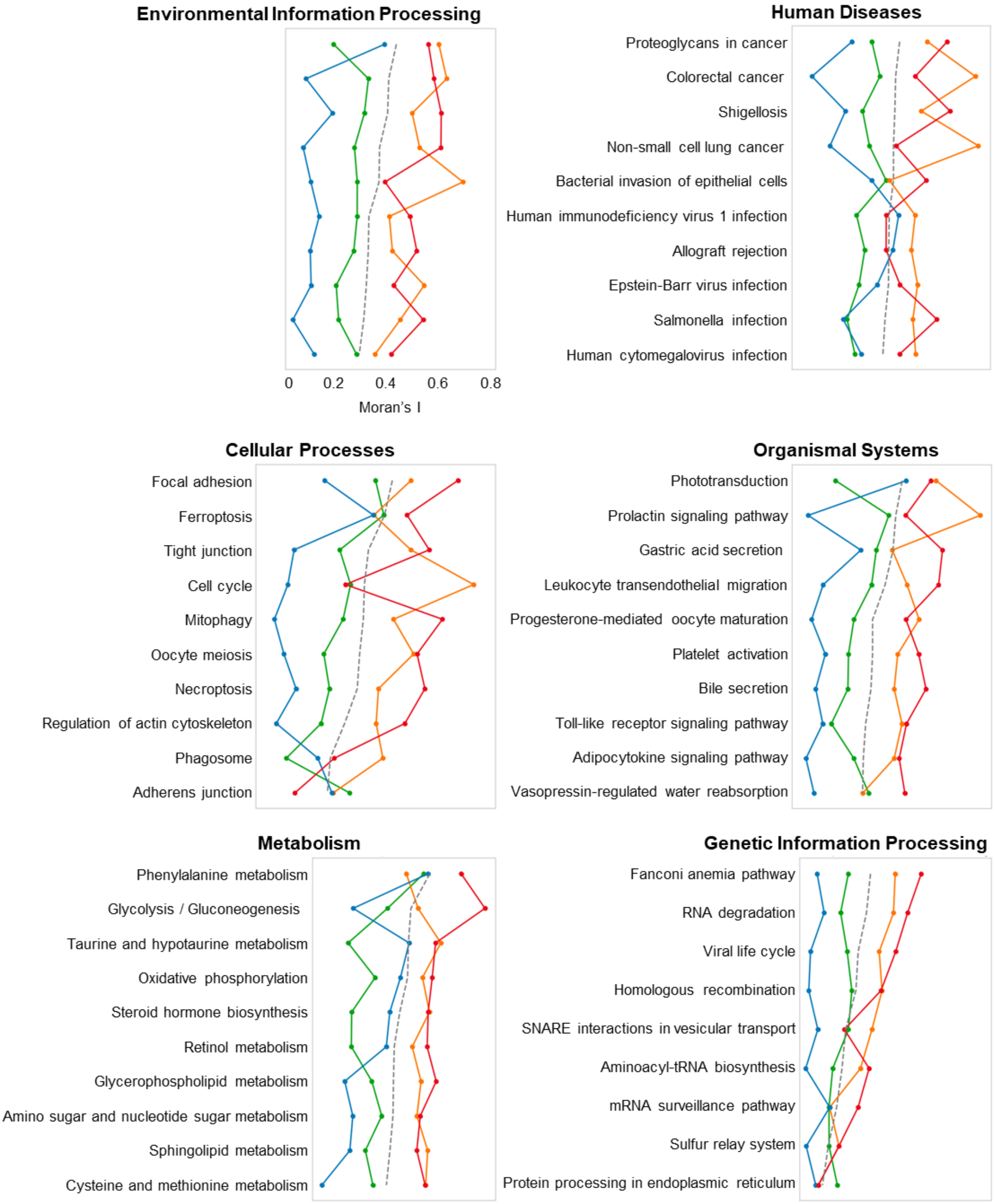


**Figure S12.** In the IDC dataset, scatterplots show the top 10 pathways by Moran's I for six pathway categories, with grey dashed lines indicating the category-specific medians.

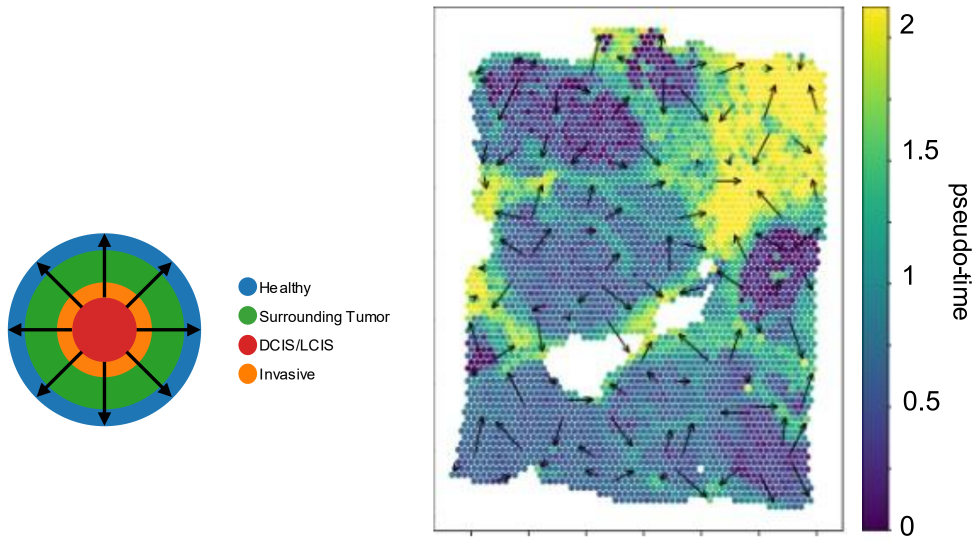


**Figure S13.** Left: The diagram, a pseudo-time trajectory map, shows the direction of this trajectory, moving from DCIS/LCIS through invasive and surrounding tumor tissues toward normal tissue. Right: Visualization of the trajectory inferred by PathCLAST, with arrows indicating the path from tissue locations with low pseudo-time to those with high pseudo-time.

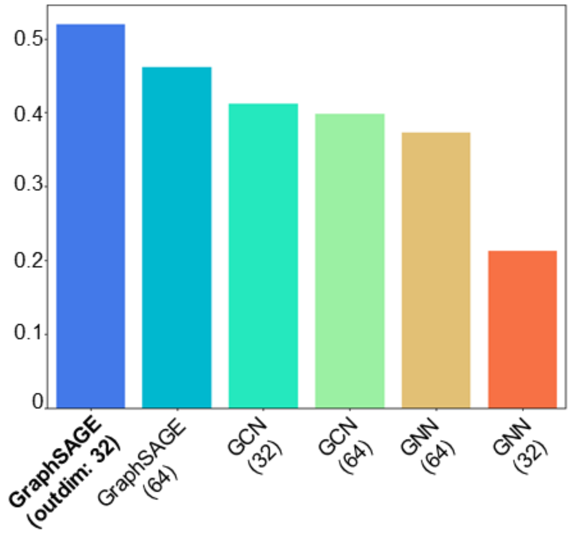


**Figure S14.** Ablation study on PathCLAST using GNN, GCN, and GraphSAGE. Performance for different pathway graph encoders and output-dimension settings was evaluated by ARI on the IDC dataset.

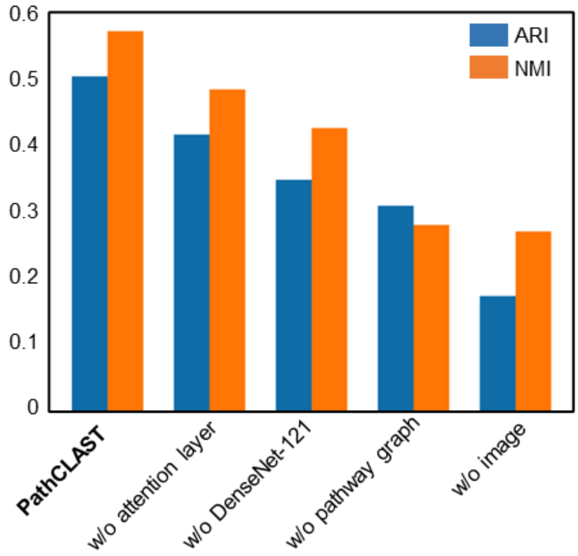


**Figure S15.** Ablation study of PathCLAST, comparing the full model with variants individually removing one module (attention layer, pretrained DenseNet – 121 weights, pathway graphs, or patch-level images). Performance was evaluated by ARI and NMI on the IDC dataset.

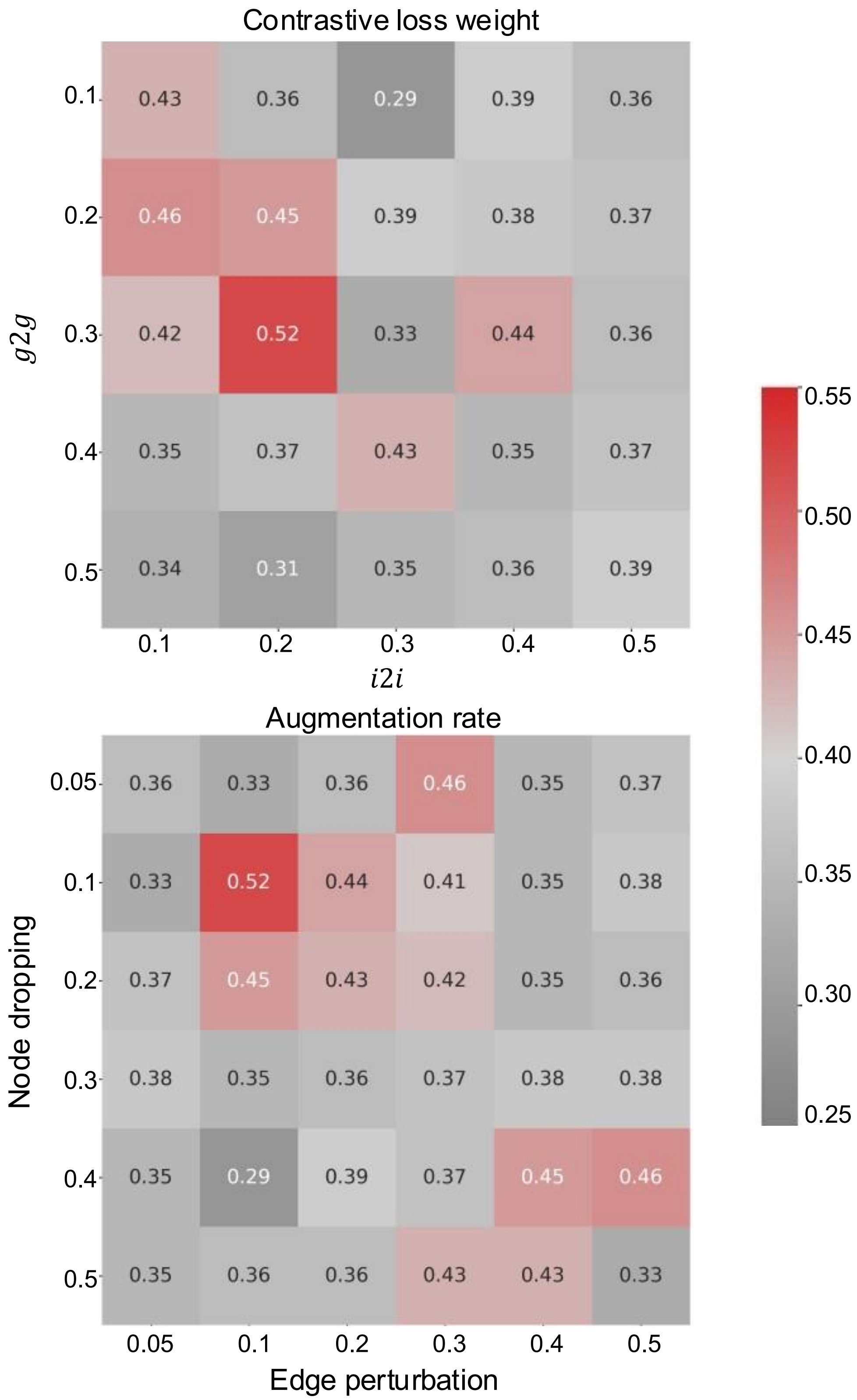


**Figure S16.** Ablation studies on the contrastive – loss weights ($L_{g2g}$ and $L_{i2i}$) and on graph – augmentations (node dropping and edge perturbation) were conducted on the IDC dataset, with performance evaluated by ARI.

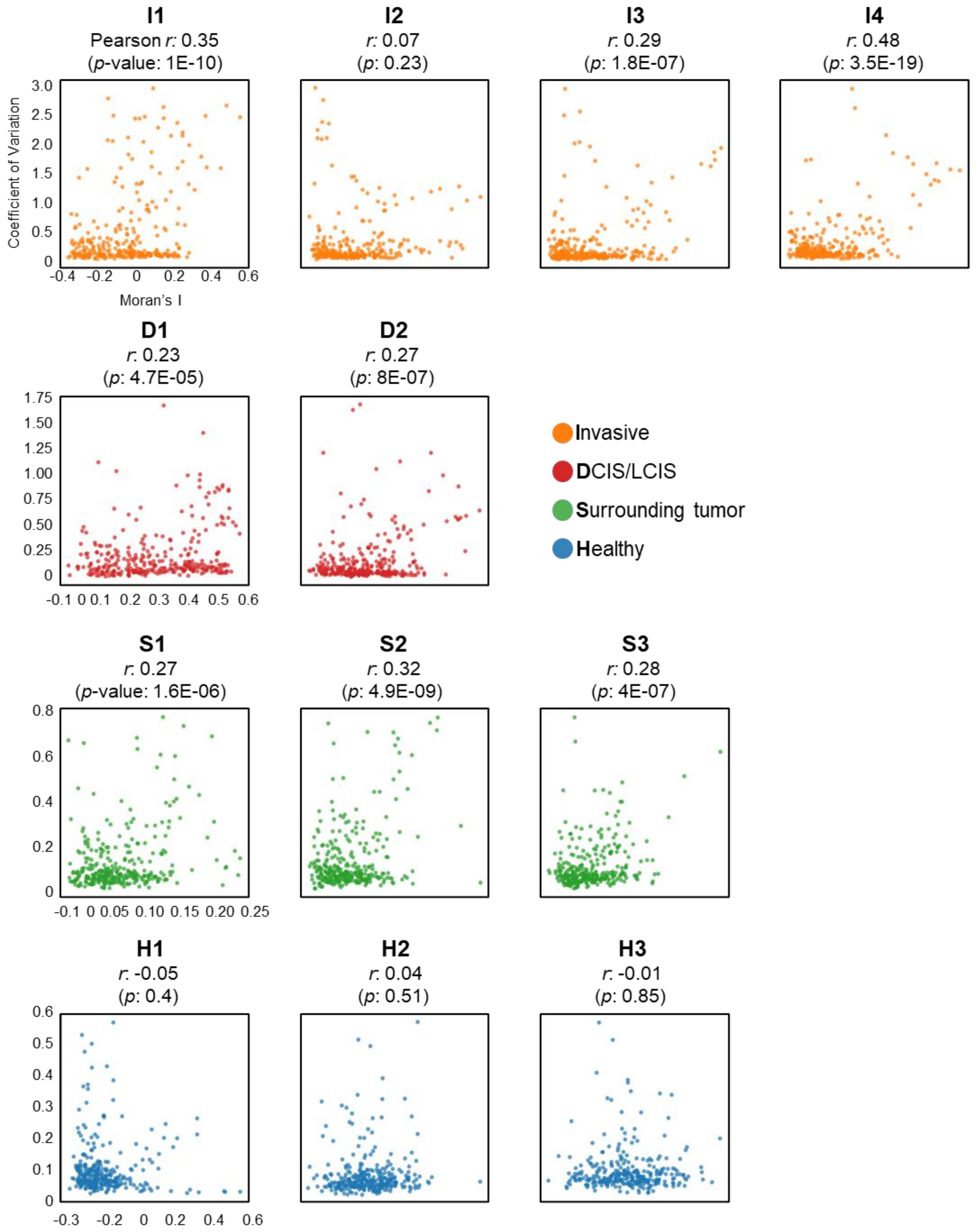


**Figure S17.** Scatterplots of Moran's I (x-axis) versus coefficient of variation (CV, y-axis) for pathways in each sub-domain of the IDC dataset.

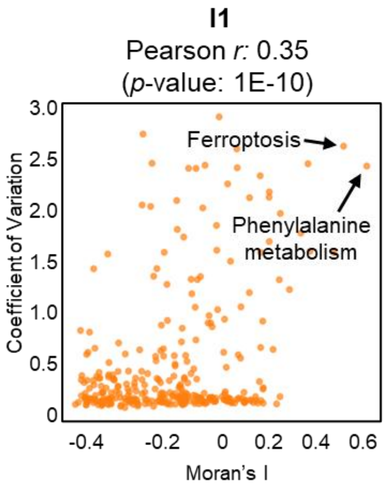


| # | Pathway Name | Moran's I | Coefficient of Variation |
|---|---|---|---|
| 1 | Phenylalanine metabolism | 0.46 | 2.36 |
| 2 | Ferroptosis | 0.42 | 2.55 |
| 3 | Prolactin signaling pathway | 0.41 | 1.53 |
| 4 | Necroptosis | 0.36 | 2.38 |
| 5 | Diabetic cardiomyopathy | 0.32 | 1.90 |
| 6 | Salmonella infection | 0.30 | 2.11 |
| 7 | Proteoglycans in cancer | 0.30 | 2.06 |
| 8 | Focal adhesion | 0.29 | 2.27 |
| 9 | Gastric acid secretion | 0.25 | 2.52 |
| 10 | Vibrio cholerae infection | 0.22 | 2.83 |

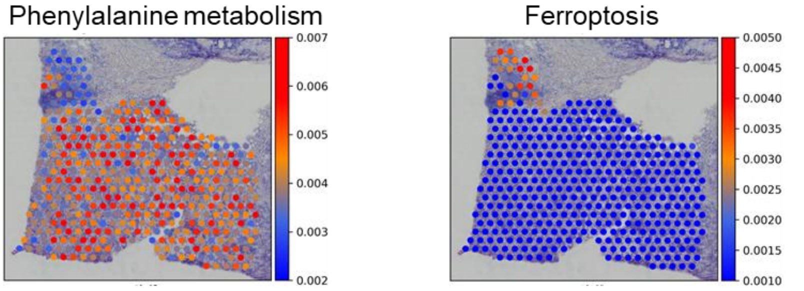


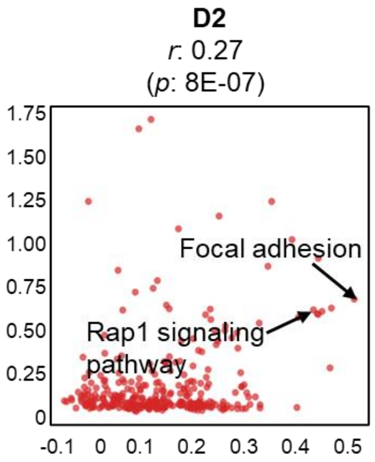


| # | Pathway Name | Moran's I | Coefficient of Variation |
|---|---|---|---|
| 1 | Focal adhesion | 0.49 | 1.25 |
| 2 | Hypertrophic cardiomyopathy | 0.45 | 1.15 |
| 3 | Dilated cardiomyopathy | 0.43 | 1.68 |
| 4 | Rap1 signaling pathway | 0.43 | 1.12 |
| 5 | Leukocyte transendothelial migration | 0.42 | 1.13 |
| 6 | Shigellosis | 0.42 | 1.09 |
| 7 | Gastric acid secretion | 0.37 | 1.88 |
| 8 | Vibrio cholerae infection | 0.36 | 2.29 |
| 9 | Fluid shear stress and atherosclerosis | 0.32 | 1.59 |
| 10 | Salmonella infection | 0.24 | 2.14 |

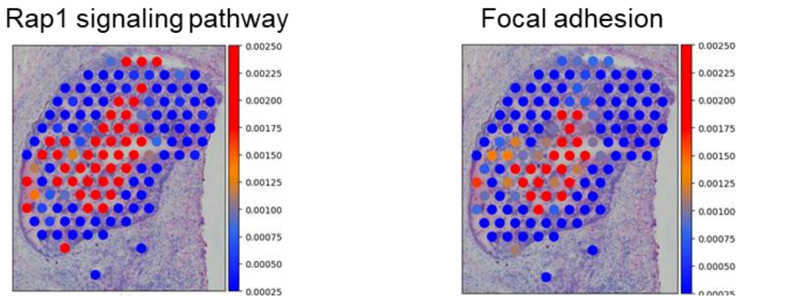


**Figure S18.** Left**:** Scatterplot showing Moran's I (x-axis) plotted against the coefficient of variation (CV, y-axis) for pathways in the indicated sub-domain. Right**:** Top 10 pathways in that sub-domain, ranked by Moran's I. Bottom**:** Spatial visualization of pathway attention weights, showing heterogeneous pathway activity across the sub-domain.
